# Longitudinal quantitative streamline tractography: robust estimation of white matter connectivity differences

**DOI:** 10.64898/2026.02.09.704742

**Authors:** Philip Pruckner, Remika Mito, David N Vaughan, Kurt G Schilling, Victoria L Morgan, Dario J Englot, Robert E Smith

## Abstract

Longitudinal probing of structural connectivity via diffusion magnetic resonance imaging (dMRI) is experiencing uptake. However, the detection of biological effects is significantly hampered by the limitations of cross-sectional streamline tractography, where even small changes in the dMRI signal can produce drastically different trajectories and therefore quantitative parameterisation; if not properly dealt with, such effects will manifest as spurious longitudinal change, which can obscure subtle biological differences. To overcome this challenge, we here introduce a novel quantitative streamline tractography framework tailored for longitudinal analysis, wherein an individual’s streamline trajectories remain fixed throughout the analysis, allowing only their ascribed density weights to vary between sessions. We present two strategies by which these quantitative streamline weights can be determined, both extensions of the widely adopted SIFT2 method. The performance of this framework is benchmarked against cross-sectional reconstruction with and without SIFT2 optimisation, in both *in silico* dMRI phantoms with known ground truths and three distinct human *in vivo* cohorts with clear a priori expectations of biological effects. We demonstrate that the proposed framework drastically reduces methodological imprecisions in synthetic dMRI phantoms and enhances statistical sensitivity and specificity to biological effects in human cohorts, enabling robust longitudinal quantification of structural connectivity.

## 1. Introduction

The human brain is commonly understood to function as a set of specialised regions embedded within a complex, interconnected network(Bassett & Sporns, 2017). Advances in diffusion magnetic resonance imaging (dMRI) based tractography have made it possible to probe the structure of this network *in vivo*, holding significant promise to improve our understanding of the human brain. In this context, longitudinal studies offer a particularly exciting opportunity, as they are highly sensitive to subtle biological effects due to the consistency of gross brain anatomy throughout the analysis, while also offering a principled basis for the inference of causal relationships from the temporal order of measurements.

The prospect of longitudinal structural connectivity analysis has been particularly advanced through the introduction of *quantitative* streamline tractography(Smith et al., 2022). These methods allow quantitative parameterisation of structural connectivity strength by enforcing correspondence between the densities of reconstructed tractograms and the fibre-densities estimated from the local diffusion model, which is typically achieved through tractogram optimisation algorithms such as SIFT2(Smith et al., 2015a). The application of these algorithms has been shown to not only improve the reproducibility of tractograms but also the correspondence with post-mortem brain dissections(Smith et al., 2015b).

Despite these advances, significant methodological variability remains when estimating longitudinal differences(Smith et al., 2015b). While tractogram optimisation methods yield quantitative connectivity information that one might intuitively expect to be more robust than non-quantitative metrics, they unfortunately remain limited by the accuracy of cross-sectional tractography, which is known to produce markedly different streamline trajectories even in scan-rescan experiments(Nath et al., 2020). Tractogram optimisation algorithms may appropriately adjust for density biases given the cross-sectionally reconstructed tractograms with which they are provided, but they are unable to account for differences in their streamline trajectories, which will ultimately manifest as spurious longitudinal change.

To increase the robustness of longitudinal neuroimaging analysis, it has proven useful to maximally leverage the shared information between timepoints, avoiding the independent reconstruction of the same gross anatomy where possible. This is well exemplified by the longitudinal FreeSurfer processing stream for anatomical image processing, which reconstructs cortical surfaces upon an unbiased within-subject template that are then adjusted based on session-specific voxel intensities(Fischl, 2012; Reuter et al., 2012; Reuter & Fischl, 2011).

The concept has since been extended to dMRI-based segmentation of major white matter bundles, using shared anatomical priors derived from an unbiased within-subject template to improve reproducibility of bundle segmentations across timepoints(Yendiki et al., 2016). Such tailored longitudinal strategies have been shown to yield more robust measurement of changes in cortical thickness and white matter microstructure than conventional cross-sectional analyses(Reuter et al., 2012; Yendiki et al., 2016).

In the domain of streamline-based tractography, the “differential tractography” method has been proposed(Yeh et al., 2019): unlike conventional fibre tracking that propagates streamlines along local fibre orientations, differential tractography specifically traces only the parts of bundles where anisotropy differences between scans exceed a certain threshold. First applications highlight the method’s potential as a biomarker for neuronal injury, showing disease-specific changes across a range of conditions(Barrios-Martinez et al., 2022; Sarica et al., 2025; Yeh et al., 2019). However, the outputs of this method are segments of white matter bundles without intrinsic quantitative properties, making them unsuitable for analysis of whole-brain structural connectivity.

In this article, we present a novel *unbiased quantitative streamline tractography* framework specifically tailored to the robust analysis of longitudinal data. Instead of reconstructing timepoints independently, we assert that an individual’s gross white matter can be represented through a single tractogram constructed in an unbiased within-subject template, thereby establishing streamline correspondence across sessions. We then optimise the densities ascribed to streamlines in the tractogram to capture the longitudinal differences in structural connectivity. We present two distinct strategies for such optimisation, both extensions of the widely adopted SIFT2 method(Smith et al., 2015a):

- The first strategy we here refer to as *symmetric* optimisation: it adopts a similar strategy to longitudinal FreeSurfer(Fischl, 2012; Reuter et al., 2012; Reuter & Fischl, 2011), first optimising per-streamline weights based on the within-subject template then adjusting them to match session-specific fibre densities.
- The second strategy we term *differential* optimisation: it shares attributes with differential tractography(Yeh et al., 2019), providing a bespoke approach to longitudinal quantitative connectivity analysis by directly optimising streamline-wise density changes to match fibre density differences between sessions.

The framework is here evaluated in the context of structural connectome analysis, quantifying longitudinal changes in Fibre Bundle Capacity (FBC)(Smith et al., 2022), a biologically meaningful connectivity metric sensitive to a bundle’s intra-axonal cross-sectional area (see Figure 1). We assess the framework’s longitudinal reconstruction error in *in silicio* dMRI phantoms with known ground truth, and its statistical sensitivity and specificity to biologically expected effects in three complementary human *in vivo* cohorts. Results are benchmarked against cross-sectional tractogram reconstruction with and without SIFT2 density optimisation.

**Fig. 1.**
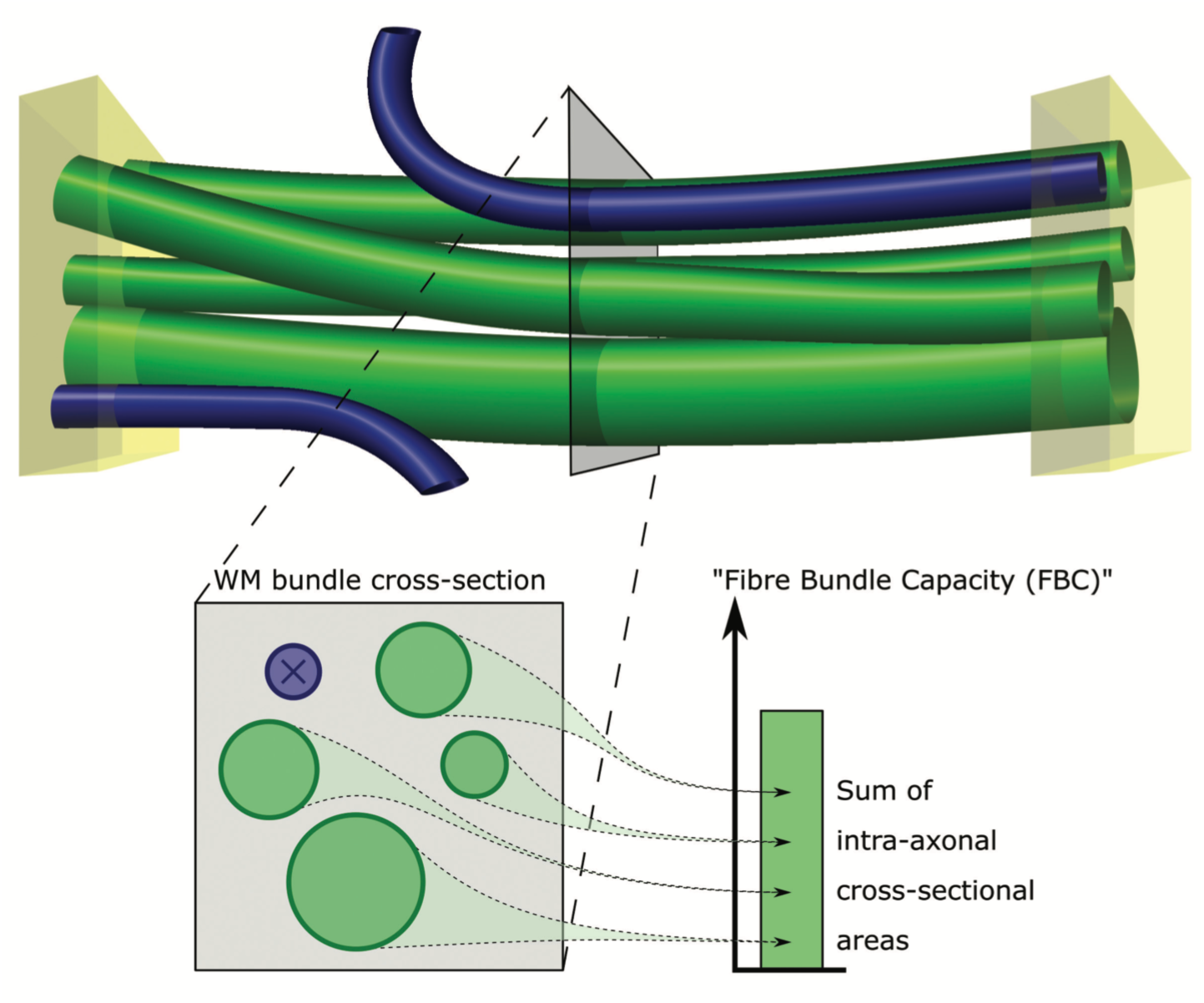
Visual depiction of the Fibre Bundle Capacity (FBC) metric. A white matter bundle of interest is defined based on its endpoints, shown as yellow cuboids. Only those fibres that are attributed to both endpoints are constituent members of that bundle (green cylinders). The FBC is defined as the sum of the intra-axonal cross-sectional areas of these fibres. Reproduced from Smith et. al(Smith et al., 2022).

## 2. Methods

The following methodological sections are separated in three parts. First, we introduce the relevant background of the proposed framework including the original SIFT2 method, the establishment of streamline correspondence across timepoints through reconstruction of an unbiased tractogram, and the novel “symmetric” and “differential” SIFT2 density optimisation strategies. The second part then outlines the data used to validate the proposed framework, including the validation in *in silicio* dMRI phantoms and human *in vivo* cohorts. The third part outlines the relevant image processing as well as the specific longitudinal connectome pipelines used for benchmarking unbiased against cross-sectional reconstruction pipelines.

To facilitate technical discussion of this article, we provided a summary of key terms and concepts in Table 1.

**Table 1.** Key terminology. Definition of key terms and concepts as used in the context of this article.

| Term | Definition |
| --- | --- |
| Fixel | Directional element within a voxel corresponding to a fibre population. |
| Fibre Density and Cross-Section | Local quantitative measurement ascribed to a fixel, representing a combination of both the microscopic intra-axonal density of fibres and the morphological modulation of such in registration to the within-subject template. For brevity here referred to as “local fibre densities”. |
| Tractogram density optimisation | The correction of density biases within streamline tractograms through calculation of appropriate streamline weights that facilitate quantitative interpretation of structural connectivity. Note that these methods are also known as “semi-global”(Smith et al., 2022) or “discriminative global” tractography(Daducci et al., 2025). |
| Streamline weight | A scalar value computed for a streamline that modulates the intra-axonal cross-sectional area ascribed to it. |
| SIFT2 | Tractogram density optimisation algorithm that determines a suitable set of streamline weights so that the density of a tractogram fits local fibre densities. |
| Unbiased (quantitative) tractogram | Whole-brain tractogram for which streamline trajectories (and streamline weights) are computed based on an unbiased within-subject template. |
| Symmetric optimisation | Density optimisation of streamline weights ascribed to an unbiased quantitative tractogram to match session-specific local fibre densities. |
| Differential optimisation | Density optimisation of streamline weights ascribed to an unbiased quantitative tractogram to directly match local fibre density differences between sessions. |
| Fibre Bundle Capacity | Inter-regional quantitative structural connectivity metric, representing the intra-axonal cross-sectional area of fibres ascribed to a given tract. |
| Number of Streamlines | Legacy structural connectivity metric with well-known biases(Jones et al., 2013), computed as the number of streamlines between regions (= streamline count). |
| Fibre count | The true number of simulated fibres connecting two regions, under direct experimental control in phantom simulations. |

### 2.1 SIFT2: original method and novel extensions

#### 2.1.1 Original method

The goal of the original SIFT2 method(Smith et al., 2015a) is to correct densities within streamline tractograms, which can arise from various biases during fibre tracking. This is achieved by determining a set of cross-sectional streamline weights that minimises the local differences between the density contribution of streamlines traversing the same fixel and local fibre densities that were estimated from the local diffusion model.

For each fixel *f*, SIFT2 determines the local tractogram density as follows:

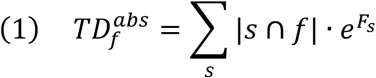

where 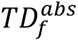 is the absolute Tractogram Density for a fixel *f,* |*s* ∩ *f*| the intersection length of streamline *s* with fixel *f,* and 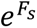 the cross-sectional weight for streamline s. Optimisation is performed on the exponential coefficient *F*_*s*_, by default initialised as *F*_*s*_= 0 ∀ s.

To enable comparison between tractogram-based densities and local fibre density estimates, the SIFT model defines the “proportionality coefficient” *μ,* which serves as global scaling between the total length of reconstructed streamlines and the total measured white matter content:

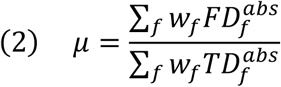

where 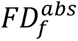 is the measured absolute Fiber Density in fixel *f*, and *w*_*f*_ the model weight for fixel *f* (which downregulates the impact of partial volume voxels, typically based on an anatomical image tissue segmentation).

The algorithm then iteratively tries to minimise the cost function:

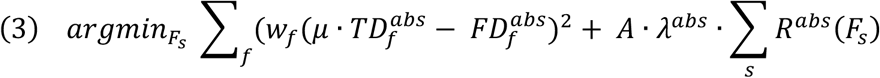

where *R*^*abs*^(*F*_*s*_) is a function computing the regularisation cost computed for streamline coefficient *F*_*s*_, ⋋^*abs*^ a user-controllable parameter for absolute regularisation strength, and *A* is a scaling parameter designed to ensure comparable effects of regularisation for different imaging and reconstruction parameters.

Finally, the Fibre Bundle Capacity(Smith et al., 2022) is computed as:

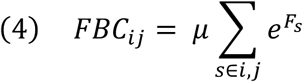

where *FBC*_*i*j_ is the Fibre Bundle Capacity for a bundle defined as connecting grey matter parcels *i* and *j*, *s* ∈ *i*, *j* selects those streamlines *s* assigned to nodes *i* and *j*, 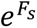 is the cross-sectional weight assigned to streamline *s*.

Detailed discussion of the regularisation term, as well as the specific choices made for this study, are in the Supplementary Materials.

#### 2.1.2 Fixing streamline trajectories throughout the analysis

One of the core premises of the proposed framework is utilisation of a common tractogram across timepoints; by precluding variations in streamline trajectories across timepoints, longitudinal changes in FBC are based only on the weights ascribed to those streamlines as driven by differences in local fibre densities (see Fig. 2). As unbalanced processing (such as utilising the tractogram reconstructed from one timepoint for the other) could introduce asymmetry biases(Reuter & Fischl, 2011), we derive such tractograms in symmetrically constructed within-subject templates. The SIFT2 algorithm is applied to determine streamline weights that achieve correspondence between the density of this tractogram and the local fibre densities computed from the template image. We here refer to such tractograms as *unbiased quantitative tractograms*, which serve as the baseline for further *symmetric* or *differential* density optimisation (see Fig. 3).

**Fig. 2.**
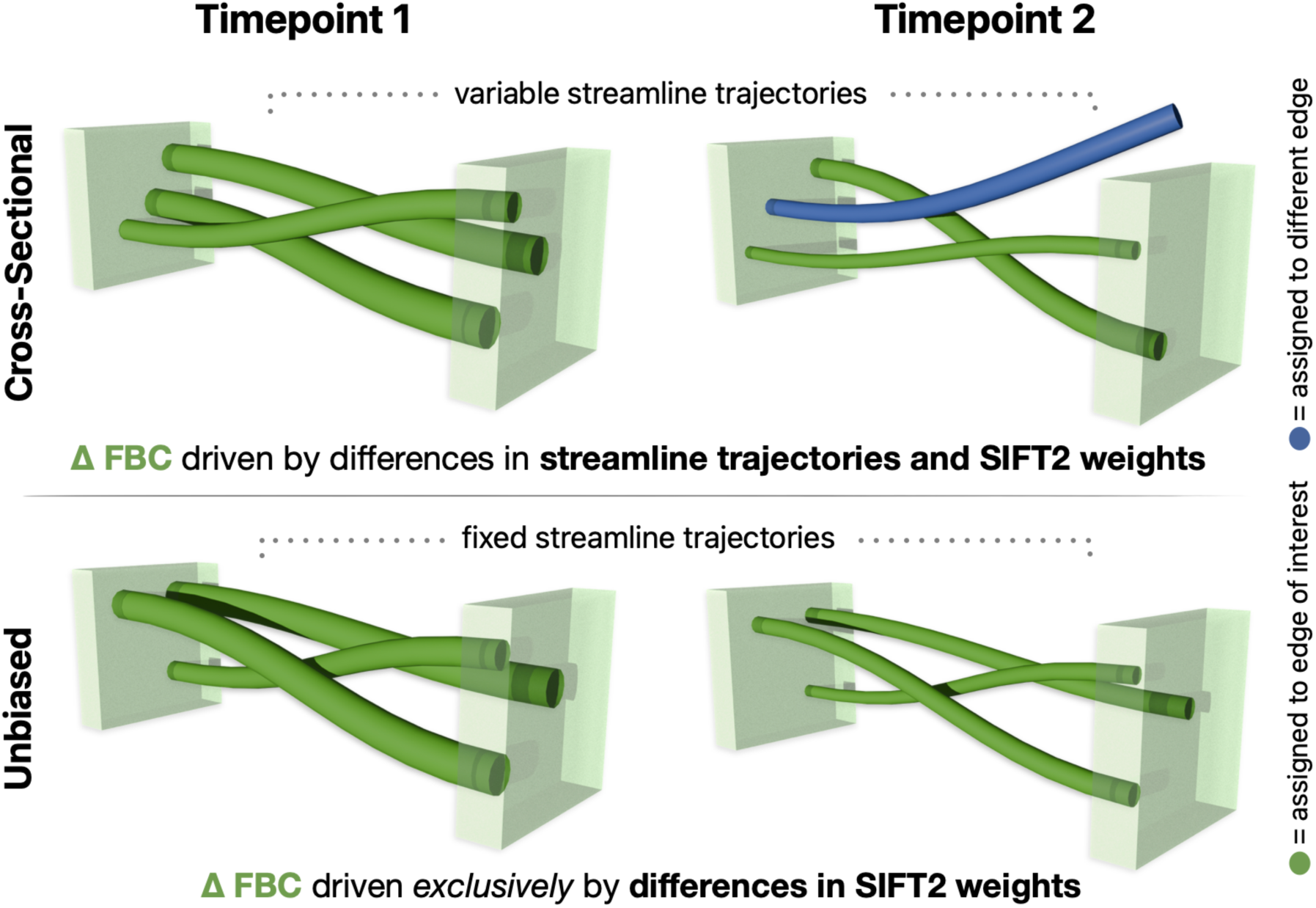
Longitudinal estimation of Fibre Bundle Capacity using cross-sectional vs. unbiased methods. Top: cross-sectional reconstruction will result in different streamline trajectories, which are not guaranteed to terminate at the same pair of regions; longitudinal FBC differences are therefore driven not only by differences in underlying fibre density, but also by differences in streamline trajectories. Bottom: unbiased reconstruction explicitly utilises the same set of streamlines across timepoints, ensuring that longitudinal FBC differences are exclusively driven by differences in their ascribed weights. Abbreviations: FBC = Fibre Bundle Capacity; SIFT = Spherical-deconvolution Informed Filtering of Tractograms.

**Fig 3.**
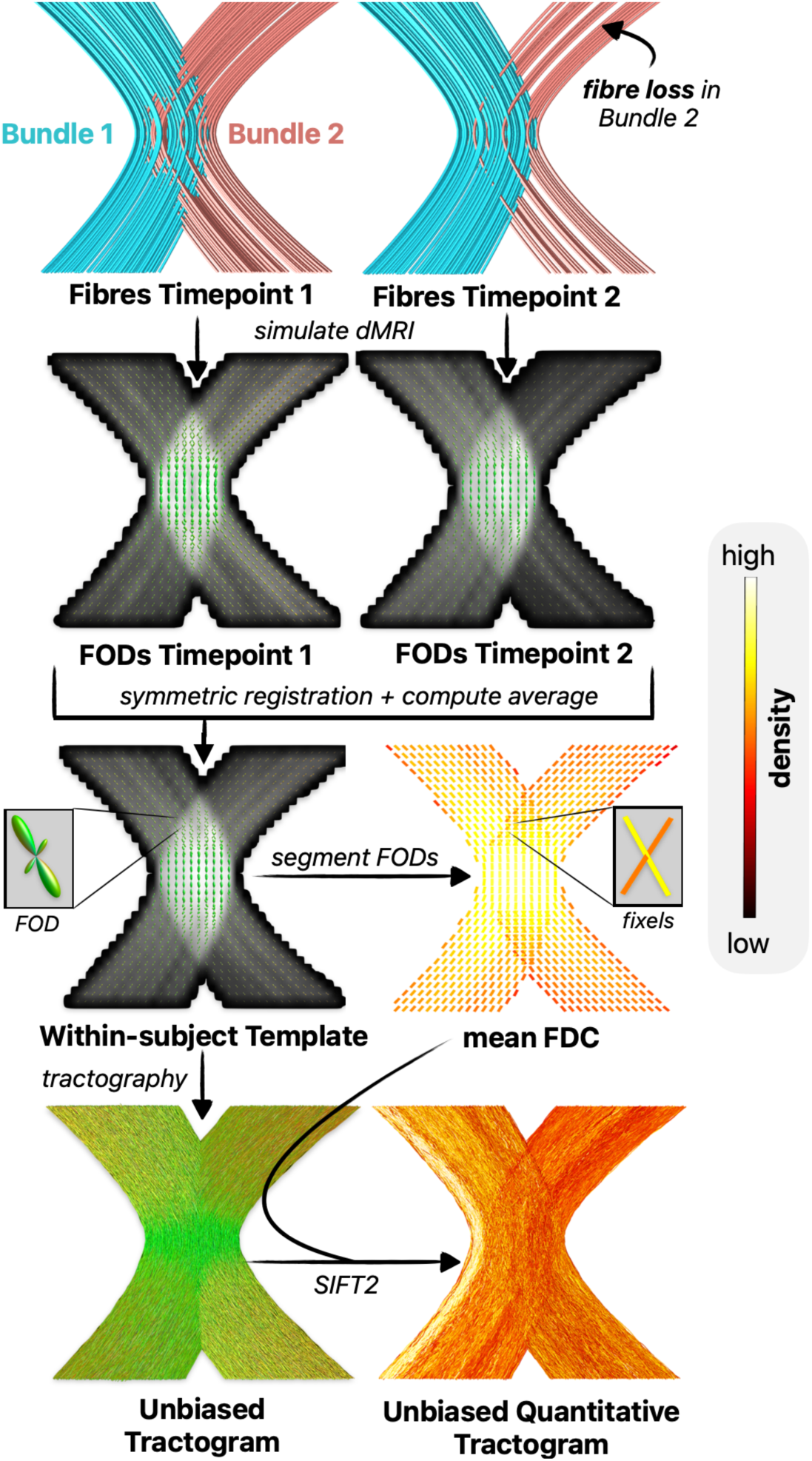
Reconstruction of unbiased quantitative tractograms. The reconstruction process is here illustrated in a synthetic phantom with simulated fibre loss in one of two kissing bundles. Image data across sessions are symmetrically aligned and averaged to produce an unbiased within-subject template. In this template, a single streamline tractogram is reconstructed. Fibre Orientation Distribution functions (FODs) are segmented into distinct directional fibre elements (fixels) to which quantitative measurements of Fibre Density and Cross-Section (FDC) are ascribed. An unbiased quantitative tractogram is constructed by optimising streamline weights based on the session-average local fibre densities. Abbreviations: SIFT = Spherical-deconvolution Informed Filtering of Tractograms.

#### 2.1.3 Symmetric SIFT2 optimisation

“Symmetric” SIFT2 optimisation follows a similar strategy to the longitudinal FreeSurfer pipeline(Fischl, 2012; Reuter et al., 2012). In this analysis stream, FreeSurfer reconstructs cortical surfaces upon an unbiased template, and then adjusts vertex placements based on session-specific voxel intensities. Symmetric tractogram optimisation can be thought of as adaptation of this strategy for quantitative streamline tractography.

Following construction of the unbiased quantitative tractogram, the SIFT2 algorithm is re-run independently per timepoint, matching the tractogram to session-specific local fibre densities, with the optimisation initialised not naively but based on the streamline weights and the proportionality coefficient *μ* within the unbiased quantitative tractogram; in this way, the resulting streamline weights will only perturb from that of the unbiased quantitative tractogram where doing so is beneficial to the quality of fit of the tractogram to those session-specific local fibre densities (see Fig. 4A).

**Fig. 4.**
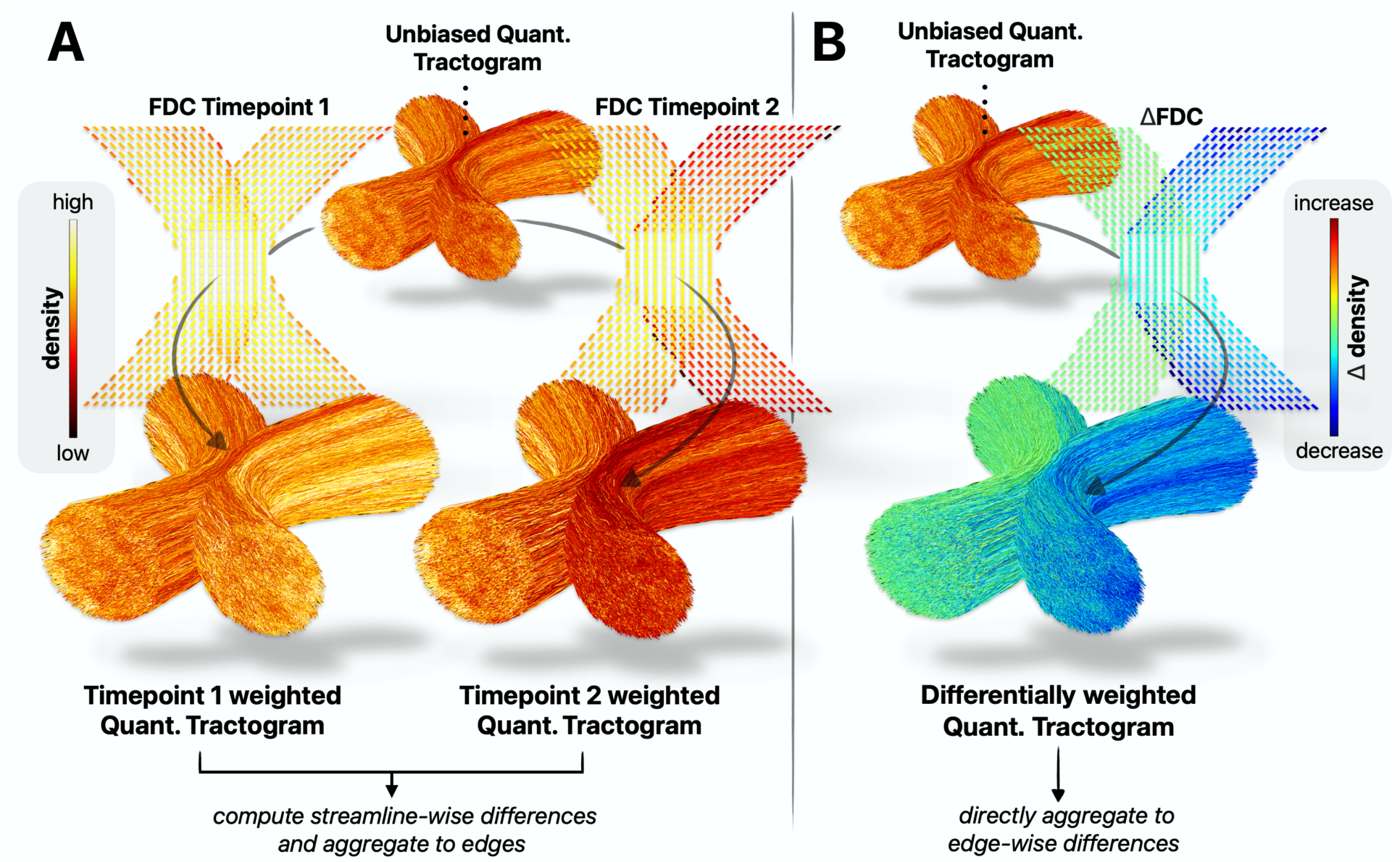
Two strategies for unbiased SIFT2 optimisation to robustly determine longitudinal structural connectivity differences. Shown are exemplar inputs and outputs of unbiased optimisation in a synthetic kissing bundles phantom with simulated fibre loss in one bundle. **A)** Symmetric optimisation initialises separate optimisation runs using the unbiased quantitative tractogram, optimising the session-average streamline weights to fit each session’s fibre-densities. **B)** Differential optimisation directly optimises the unbiased quantitative tractogram to fit inter-session fibre-density differences. Abbreviations: FDC = Fibre Density and Cross-section, SIFT = Spherical-deconvolution Informed Filtering of Tractograms.

Following optimisation of the streamline weights independently per timepoint, the change in FBC for a given tract is then computed as the sum of the differences in streamline weights between the two time points for all streamlines assigned to that tract:

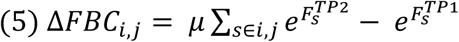

where Δ*FBC*_*i*,j_ is the longitudinal change in Fibre Bundle Capacity between parcels *i* and *j*, *μ* is the proportionality coefficient of the unbiased quantitative tractogram, and 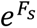 is the cross-sectional weight of streamline *s* at either the first or second timepoint.

The utility of this approach has been demonstrated in a first application study(Pruckner et al., 2025).

#### 2.1.4 Differential SIFT2 optimisation

Differential SIFT2 optimisation shares attributes with the “differential tractography” method(Yeh et al., 2019), which performs fibre tracking on inter-session differences to reconstruct the trajectories of only those segments of bundles that exhibit a longitudinal change above a pre-defined threshold. Differential optimisation extends this idea to the domain of quantitative streamline tractography, optimising the streamline weights within an unbiased whole-brain tractogram to fit such inter-session differences (see Fig 4B).

For each fixel *f*, differential optimisation defines the local *differential* tractogram density as follows:

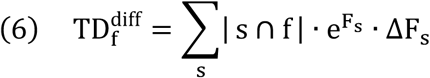

where 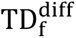 is the *differential* Tractogram Density for fixel *f*, |s ∩ f| the intersection length of streamline *s* with fixel *f,* 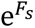 the precomputed density weight for streamline *s* within the unbiased quantitative tractogram, and ΔF_s_is the so-called “delta coefficient” for streamline*s* (note that this is a *unique* variable, not to be read as change in *F*_*s*_). Differential optimisation optimises these delta coefficients ΔF_s_, determining the relative modulation of the precomputed unbiased weights necessary to fit local fibre density differences between sessions.

The algorithm iteratively tries to minimise the cost function:

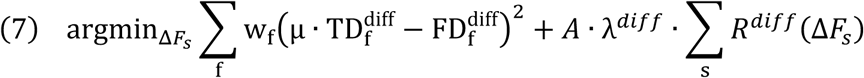

where 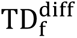 the differential Track Density for fixel *f*, 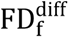 is the fibre density half-difference between timepoints for fixel *f*, λ^*diff*^ a user-controllable parameter for differential regularisation strength, and *R*_*diff*_(*ΔF*_*s*_) a regularisation function applied to the delta coefficient of streamline *s*.

The synergy between the use of mean fibre densities across timepoints for the derivation of the unbiased quantitative tractogram, the prior definition of *ΔF*_*s*_, and the utilisation of a half-difference for 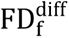, is that the most extreme scenario possible, where a bundle is present in one timepoint and absent in the other, would manifest as *ΔF*_*s*_ = ±1.0, defining an intuitive range within which to constrain these values (see Supplementary Methods).

The longitudinal FBC difference between timepoints is then computed as:

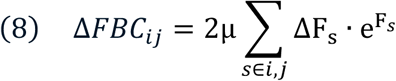

where Δ*FBC*_*i*j_ is the longitudinal change in Fibre Bundle Capacity for the bundle defined based on connectivity between grey matter parcels *i* and *j*, *s* ∈ *i*, *j* a streamline *s* assigned to nodes *i* and *j*, and ΔF_s_ · 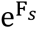 the so-called “delta weight” encoding the absolute contribution of streamline *s* to the observed local fibre density difference signal.

Discussion of the differential regularisation term, as well as the specific choices selected for this study, are detailed in the Supplementary Materials.

### 2.2 Validation

#### 2.2.1 Validation in *in silicio* dMRI phantoms

Synthetic dMRI phantoms were employed to quantify the accuracy of longitudinal connectivity estimates obtained through connectome pipelines later described in this section. Simulations were based on the previously published DiSCo3 phantom(Rafael-Patino et al., 2021), a composition of fibres that mimic realistic anatomical white matter complexities. We introduced three distinct effects into this phantom by manipulating the fibre count of specific bundles between timepoints, creating three longitudinal models (see Fig 5A):

- **Model 1 – Single Bundle**: a 75% fibre count decrease in a single isolated bundle.
- **Model 2 – Crossing Bundles**: changes introduced into two bundles that cross one another; one with a 50% fibre count decrease, the other with a 100% increase.
- **Model 3 – Central Lesion**: a volumetric lesion was placed in the centre of the phantom, with all fibres intersecting the lesion removed from the second timepoint. This mimics complex lesion-related fibre count decreases of different magnitudes in a range of bundles.

**Fig. 5.**
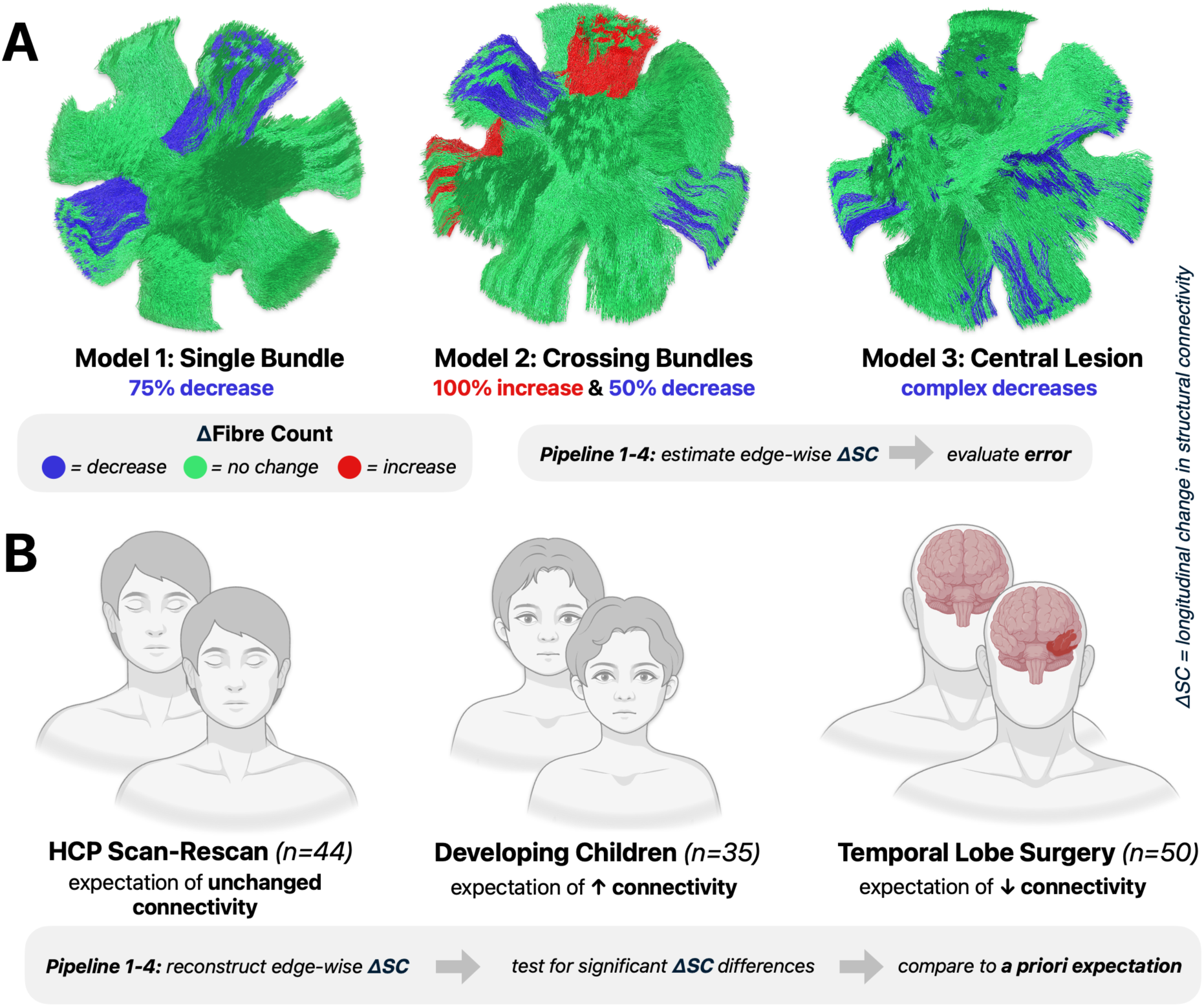
Simulated and expected longitudinal effects in synthetic dMRI phantoms and human *in vivo* data. **A)** Three synthetic models for quantitative validation of connectome pipelines based on the DiSCo3 phantom, from which artificial dMRI images were synthesised. Estimated changes in structural connectivity compared to the ground truth fibre count changes. B) *In vivo* validation comprised three independent datasets, with different a priori expectations of effects against which statistically significant changes were compared. *Created in BioRender. Pruckner, P. (2026)* https://BioRender.com/dy5zkqh

Based on these models, we synthesised artificial dMRI images using Fiberfox(Neher et al., 2014). Tissue components were simulated using a ball and stick model, with tissue properties refined to yield diffusion-weighted images that approximated empirical *in vivo* human data. For diffusion gradient encoding, we used a subset of the shells provided in the gradient table of the original dataset(Rafael-Patino et al., 2021) (*b*=0, 1000, 2000, 3000). Furthermore, we added Rician noise(Garyfallidis et al., 2014) at a signal-to-noise ratio of 20, mimicking realistic *in vivo* acquisitions. Further information about the simulation parameters and refinement of tissue properties is provided in the Supplementary Materials.

To facilitate comparison of tractography-based connectivity measures against the known ground truth fibres, all estimated changes were scaled to the true fibre count using a global scaling factor:

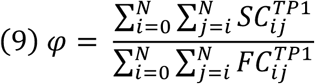

where 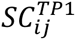 the structural connectivity between parcels *i* and *j* at the first timepoint, and 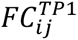 the ground truth fibre count between parcels *i* and *j* at the first timepoint (computed from phantom fibres used to simulate dMRI acquisitions).

For each longitudinal connectome pipeline, we quantified the bundle-wise longitudinal error, i.e. the absolute deviation of structural connectivity changes from the ground truth fibre count change:

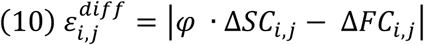

where *φ* · Δ *SC*_*i*,j_ is the globally scaled structural connectivity change between parcels *i* and *j,* Δ*FC*_*i*,j_ the true fibre count change between parcels *i* and *j*.

Statistical differences in bundle-wise absolute error distributions were assessed between pipelines through the non-parametric Kruskal-Wallis test (a non-parametric test was chosen as the empirical distribution of absolute errors (bounded [0, infinity]) violate Gaussian normality assumptions). If an overall statistical difference was observed, we conducted pairwise Dunn’s post-hoc tests with Holm correction for multiple comparisons to determine the specific pipeline pairs driving this result. Statistical tests were considered as significant at α=0.05.

#### 2.2.2 Validation in human *in vivo* cohorts

For *in vivo* validation of connectome pipelines, we chose three complementary cohorts with distinct expectations of biological effects (see Fig 5B):

- **Human Connectome Project (HCP) Scan-Rescan**(Essen et al., 2013) (n=44): expectation of no difference in structural connectivity.
- **Developing Children**(Leon Y Cai et al., 2021) (n=35): expectation of widespread increases in structural connectivity.
- **Temporal Lobe Surgery** (n=52): expectation of pronounced structural connectivity decreases ipsilateral and proximal to resection.

Baseline information of each cohort is provided in Table 2, with further cohort details and information on MRI sequences provided in the Supplementary Materials.

**Table 2.** Baseline information of *in vivo* cohorts. Abbreviations: ATL = anterior temporal lobectomy, EPI = echo-planar imaging, LITT = laser interstitial thermal therapy, MPRAGE = Magnetization-Prepared Rapid Gradient-Echo, SAHE = selective amygdalohippocampectomy, TA = time of acquisition, TFE = turbo field echo.

|  | **HCP Scan-Rescan** | **Developing Children** | **Temp. Lobe Surgery** |
| --- | --- | --- | --- |
| **Cohort** | 44 healthy adults | 35 healthy children | 52 adult epilepsy patients |
| **Age Range** | 22-35 years | 6-8 years | 18-68 years |
| **Scan Interval** | 6 months | 1 year | 19 months |
| **Intervention** | N/A | N/A | SAHE (n=36)<br>ATL (n=13)<br>LITT (n=3) |
| **Scanner** | Siemens 3T<br>Connectom Skyra | Philips 3T Achieva | Philips 3T Achieva |
| **dMRI acquisition** | spin-echo EPI sequence with $b=0$ (18), $b=1000$ (90), $b=2000$ (90), $b=3000$ (90) at $2.5\text{mm}^3$ isotropic. TA = ~10 min. | spin-echo EPI sequence with $b=0$ (16), $b=1000$ (40), $b=2000$ (56) at $2.1 \times 2.1 \times 2.2\text{mm}^3$. TA ~10 min. | spin-echo EPI sequence with $b=0$ (1), $b=1600$ (96) at $2.5\text{mm}^3$. TA ~11 min. |
| **T1-weighted acquisition** | 3D MPRAGE at $0.7\text{mm}^3$. Acquisition time = ~8 min. | 3D T1TFE at $1\text{mm}^3$. Acquisition time = ~4 min. | 3D T1TFE at $1\text{mm}^3$. Acquisition time = ~2 min. |

For each cohort and pipeline, we investigated the sensitivity and specificity in statistical detection of biologically expected effects. Statistical inference was performed using the Threshold-Free Network Based Statistic (TFNBS) method(Baggio et al., 2018), an extension of the widely-adopted Network Based Statistic(Zalesky et al., 2010) framework that obviates the need for a user-defined cluster forming threshold by integrating across the full range of possible thresholds to enhance connected subnetworks that show consistent edge-wise effects (enhancement parameters were set to *E* = 0.4 and *H* = 3 based on prior work(Vinokur et al., 2015a, 2015b)). For each pipeline and cohort, we counted separately the number of edges with statistically significant increases versus decreases and compared the prevalence and locations of these changes to a priori expectations.

For statistical testing, we used the following null models:

- HCP Scan-Rescan: 0 ∼ *β*_0_ · Δ*SC*

, where Δ*SC* is the edge-wise longitudinal change in structural connectivity.

- Developmental: 0 ∼ *β*_0_ · Δ*SC* + *β*_1_ · Δ*Motion*

, where Δ*SC* is the edge-wise longitudinal change in structural connectivity and Δ*Motion* is the difference between sessions of the average relative frame-wise displacement as estimated during preprocessing(Andersson & Sotiropoulos, 2016). Note that intracranial volume was not included as covariate, given we were interested in detection of maturation in the most general sense (as opposed to effects over and above global brain growth).

- Temporal Lobe Surgery: 0 ∼ *β*_0_ · Δ*SC* + *β*_1_ · *Intervention*

, where Δ*SC* is the edge-wise longitudinal change in structural connectivity, and Intervention the surgical approach performed (i.e. either anterior temporal lobectomy, selective amygdalohippocampectomy or laser interstitial thermal therapy; see Table 2).

Non-parametric shuffling for null distribution generation was performed through sign-flipping of model residuals(Winkler et al., 2014), with separate testing of hypotheses *β*_0_ > 0 and *β*_0_ < 0 and significance threshold α=0.05.

### 2.3 Processing of image data

#### 2.3.1 Preprocessing

Preprocessing for *in vivo* data was performed as follows:

- HCP Scan-Rescan data were preprocessed through the minimal HCP preprocessing pipeline(Glasser et al., 2013), including *b*=0 intensity normalisation, motion and eddy current correction and EPI distortion correction(Andersson et al., 2003; Andersson & Sotiropoulos, 2016) (FSL(Jenkinson et al., 2012)), geometric gradient nonlinearity distortion correction(Bammer et al., 2003; Jovicich et al., 2006), and registration to the T1-weighted image.
- Developing Children and Temporal Lobe Surgery data were preprocessed using PreQual(Leon Y. Cai et al., 2021), a standardised dMRI preprocessing pipeline. In short, this included Marchenko-Pastur PCA Denoising(Cordero-Grande et al., 2019; Veraart et al., 2016) (MRtrix3(Tournier et al., 2019)), inter-scan intensity normalisation, motion and eddy current correction and EPI distortion correction(Andersson et al., 2003; Andersson & Sotiropoulos, 2016) (FSL(Jenkinson et al., 2012)) using a synthetic *b*=0 image generated from the T1-weighted image (SyNb0-DisCo)(Schilling et al., 2019).

All *in vivo* T1-weighted images were bias field corrected(J. G. Sled et al., 1998), intensity normalised(Dale et al., 1999) and rigidly registered to respective dMRI images.

#### 2.3.2 Estimation of tissue-specific ODFs

For each phantom and subject, we estimated tissue response functions of white matter, grey matter and cerebrospinal fluid using a data-driven process(Dhollander et al., 2016, 2019). Tissue response functions of *in vivo* subjects were averaged within each cohort, a prerequisite for inter- and intra-subject comparison of derived quantitative metrics(Raffelt, J.-Donald Tournier, et al., 2012; Smith et al., 2022). Local orientation distribution functions (ODFs) for each tissue type were derived using multi-shell multi-tissue constrained spherical deconvolution where applicable (Scan-Rescan, Developing Children and phantoms); single-shell three-tissue constrained spherical deconvolution(Dhollander et al., 2016) was used for the Temporal Lobe Surgery cohort due to use of a single-shell acquisition.

#### 2.3.3 Within-subject template generation

For unbiased reconstruction pipelines, we aggregated longitudinal data of each phantom/subject to produce within-subject templates:

- For DiSCo3 phantoms, synthetic dMRI images were generated on a common voxel grid, so that the session-wise white matter ODFs were simply averaged to produce unbiased white matter ODF templates.
- *In vivo* sessions were first symmetrically aligned using T1-weighted image information. Rigid registration was performed using FreeSurfer’s robust template registration(Reuter et al., 2012). For subjects where morphological differences are expected between scans (Developing Children, Temporal Lobe Surgery), images were additionally non-linearly refined using ANTs’ symmetric-normalization diffeomorphic registration(Avants et al., 2011). The obtained transforms were applied to both T1-weighted images and white matter ODFs (including modulation(Raffelt, J.-Donald Tournier, et al., 2012) and reorientation(Raffelt, J-Donald Tournier, et al., 2012)), and these were averaged to produce unbiased T1-weighted and white matter ODF within-subject templates.

#### 2.3.4 Longitudinal FreeSurfer

To obtain parcellations for *in vivo* subjects, we reconstructed T1-weighted within-subject templates as per longitudinal FreeSurfer pipeline(Fischl, 2012; Reuter et al., 2012) or a modification thereof(Hoffmann et al., 2020; Pruckner et al., 2025):

- For subjects where the within-subject template was generated exclusively using rigid registration (HCP Scan-Rescan), the default longitudinal FreeSurfer pipeline was run, which uses the same registration strategy(Reuter et al., 2012) as was used for prior within-subject template generation.
- For subjects where the within-subject template was generated using rigid registration with additional non-linear refinement (Developing Children, Temporal Lobe Surgery), we interrupted the base template reconstruction after its initial stage, and manually injected the non-linear T1-weighted template so that it is reconstructed instead of the default rigid template(Hoffmann et al., 2020; Pruckner et al., 2025).

By ensuring that both longitudinal FreeSurfer(Fischl, 2012; Reuter et al., 2012) as well as the here proposed unbiased quantitative tractography pipelines reconstruct anatomy from the same/spatially aligned templates, all outputs intrinsically live in a common space. This is beneficial for downstream connectome construction, where endpoints of streamlines produced from the white matter ODF template are assigned to cortical parcels derived from the T1-weighted template. Note that this common output space can be further leveraged to comprehensively study neuroanatomical change, e.g. assessing correspondence of structural connectivity changes with changes in cortical thickness(Pruckner et al., 2025).

#### 2.3.5 Pipelines 1 & 2: cross-sectional reconstruction of streamline count and FBC

For these two pipelines, longitudinal connectivity changes were reconstructed strictly cross-sectionally. From individual session data, we derived tissue segmentations (phantoms: as provided in the original dataset(Rafael-Patino et al., 2021), *in vivo:* from T1-weighted images(Smith et al., 2020)) for Anatomically Constrained Tractography(Smith et al., 2012) (ACT), as well as parcellations (phantoms: as provided in the original dataset(Rafael-Patino et al., 2021); *in vivo*: Desikan Killiany(Desikan et al., 2006) + subcortical regions). We generated session-specific tractograms (iFOD2(Tournier et al., 2010), ACT(Smith et al., 2012)) from individual session white matter ODFs (phantoms: 20k streamlines; in vivo: 10M streamlines).

The key distinction between Pipeline 1 and Pipeline 2 is the metric of structural connectivity quantified per connectome edge:

- Pipeline 1 quantified inter-regional connectivity as the Number of Streamlines (NoS) connecting pairs of regions. Differential connectomes were computed as the differences in these counts between timepoints for each edge. To facilitate comparison with other pipelines, longitudinal changes of in vivo data were scaled to an equivalent range of FBC differences through multiplication by the proportionality coefficient *µ* of the first session (this scaling was not performed for *in silicio* validation, as comparison between pipelines was facilitated through normalisation to the underlying fibre count).
- Pipeline 2 quantified inter-regional connectivity as the Fibre Bundle Capacity (FBC) measure, computed through optimisation of streamline densities using the SIFT2 method(Smith et al., 2015a). Fixel-wise measures of Fibre Density and Cross-Section (FDC)(Raffelt et al., 2017a) were calculated from each session’s white matter ODFs and their transforms to the within-subject template space(Raffelt et al., 2017a). Differential connectomes were computed by quantifying the difference in FBC between timepoints for each edge.

A schematic overview of Pipelines 1 and 2 is shown in Fig 6A.

**Fig. 6.**
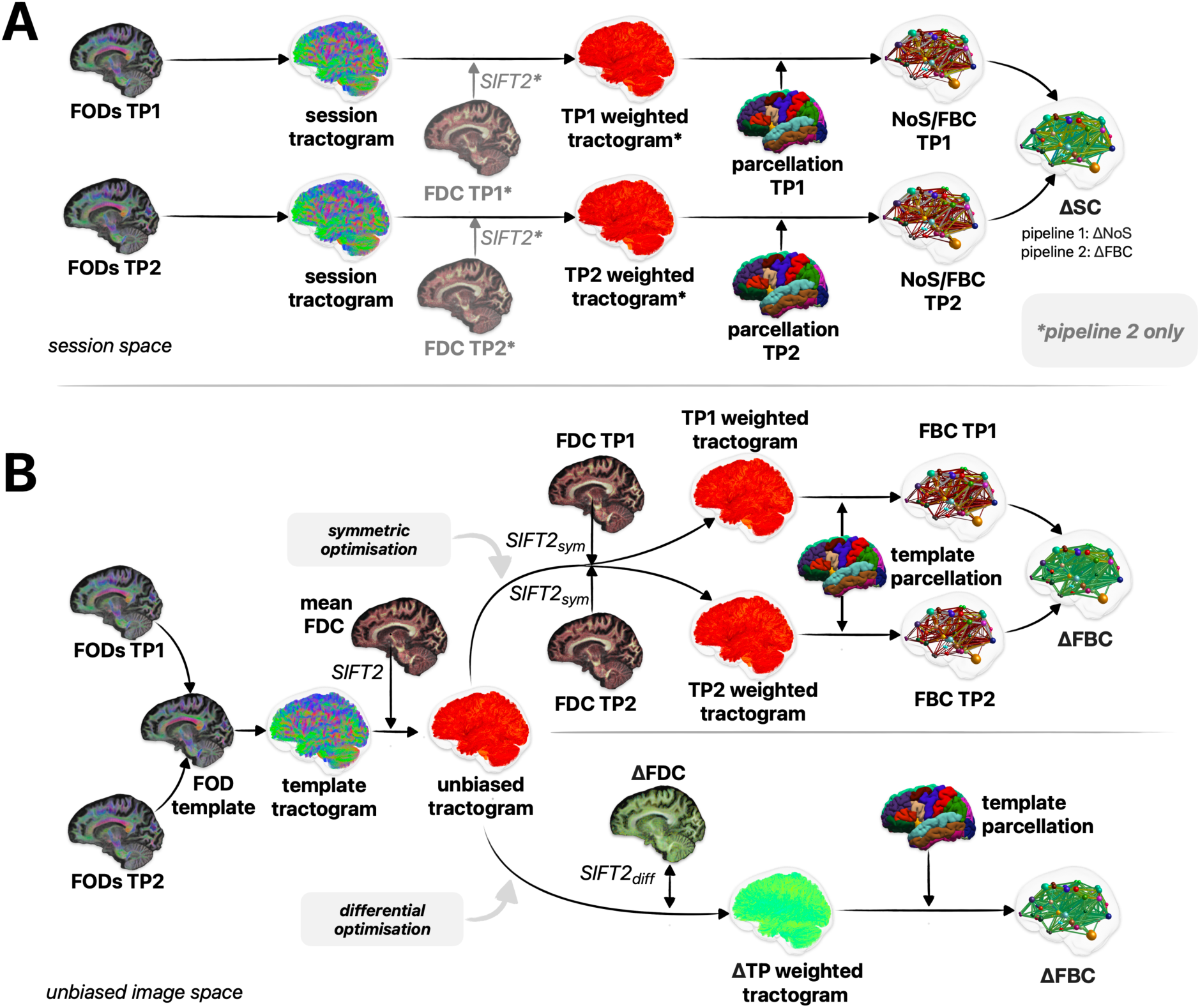
Overview of employed longitudinal connectome pipelines. Shown are the key processing steps involved in each pipeline to assess longitudinal connectivity differences at the connectome level. **A)** Cross-sectional pipelines. Session-wise tractograms and connectomes are derived independently, and from these connectomes longitudinal differences are computed. The key distinction between pipeline 1 (NoS) and pipeline 2 (cross-sectional FBC) is the application of tractogram density optimisation in the latter, as indicated by asterisks/reduced opacity. **B)** Unbiased pipelines. For each subject, a single unbiased quantitative tractogram is constructed from a within-subject template. Pipeline 3 (symmetric optimisation) uses this tractogram to initialise refinement of streamline weights to match local fibre density measurements at each timepoint, with longitudinal differences calculated from the resulting session-wise connectomes. Pipeline 4 directly optimises streamline differential weights for this tractogram to match pre-computed fibre density differences between sessions, which can be directly aggregated as a differential connectome. Diff = differential FBC = Fibre Bundle Capacity, FDC = Fibre Density and Cross-Section, FOD = Fibre Orientation Distribution Function, NoS = number of streamlines, SIFT = Spherical-deconvolution Informed Filtering of Tractograms, sym = symmetric, TP = timepoint, ΔTP = longitudinal difference between timepoints.

#### 2.3.6 Pipelines 3 & 4: symmetric and differential reconstruction of FBC

Within per-subject templates, we derived tissue segmentations (phantoms: as provided in the original dataset(Rafael-Patino et al., 2021), *in vivo:* from T1-weighted templates(Smith et al., 2020)) for ACT(Smith et al., 2012) as well as parcellations (phantoms: as provided in the original dataset(Rafael-Patino et al., 2021); *in vivo*: Desikan Killiany(Desikan et al., 2006) + subcortical regions). FOD segmentation to yield fixels(Smith et al., 2013) was performed independently for both the subject ODF templates and the session-specific ODFs transformed to those templates. The Fibre Density metric was computed for session data(Raffelt et al., 2017a) and projected to subject template fixels using an advanced fixel correspondence algorithm to mitigate artifactual longitudinal differences due to changes in FOD shape(Smith & Connelly, 2018). The Fibre Cross-section metric(Raffelt et al., 2017a) was computed for template fixels and multiplied by corresponding Fibre Density values to derive the combined metric of FDC. For each template fixel, the mean and longitudinal change in FDC across timepoints was computed. Unbiased template tractograms were generated by performing tractography upon the subject FOD template (iFOD2(Tournier et al., 2010), ACT(Smith et al., 2012); phantoms: 20k streamlines; in vivo: 10M streamlines). Quantitative streamline weights were derived using SIFT2(Smith et al., 2015a), using mean FDC across sessions as target local fibre densities.

The key distinction between Pipeline 3 and Pipeline 4 is the subsequent computation for generation of differential connectomes:

- Pipeline 3 involves additional execution of the original SIFT2 algorithm for each session, using: 1) streamlines from the unbiased quantitative tractogram; 2) session-specific FDC values projected to subject template fixels as the target local fibre densities; 3) initial values for quantitative streamline weights as computed by the prior execution of SIFT2 where the target local fibre densities were the mean FDC values across sessions. From these, structural connectomes based on the FBC metric were computed per session, with their difference forming the differential connectome.
- Pipeline 4 uses the novel augmentation of the SIFT2 method tailored to differential analysis, computing streamline differential weights for the unbiased quantitative tractogram that fit the pre-computed fixel-wise FDC difference. Aggregation of the resulting data for streamlines ascribed to each connectome edge directly yields the differential connectome.

A schematic overview of Pipelines 3 and 4 is shown in Fig 6B. Additional steps implemented to account for gross tissue resections in surgical cases can be found in the Supplementary Materials.

## 3 Results

### 3.1 Pipeline performance in *in silicio* dMRI phantoms

Across all evaluated models, unbiased reconstruction pipelines yielded significantly lower bundle-wise errors (*ε*^*diff*^) compared to cross-sectional pipelines (p<10^-4^). Only minimal (statistically insignificant) discrepancy was found between “symmetric” and “differential” unbiased reconstruction pipelines.

### 3.2 Statistical sensitivity and specificity to expected *in vivo* effects

We evaluated each pipeline’s ability to detect biologically expected effects, testing for significantly different edges (see Figure 8). Results were as follows:

- In HCP Scan-Rescan data, Pipelines 1 & 2 showed a single edge with an effect small in magnitude but nevertheless statistically significant between the left parahippocampal gyrus and the left putamen. Pipelines 3 & 4 found several changes (increases/decreases: 17/40 [symmetric optimisation] & 12/33 [differential optimisation]); effect sizes were generally small in magnitude, except for a moderate increase within the edge connecting the anterior rostral frontal gyri (note that this was a non-artefactual finding that is examined in detail in the Discussion).
- In Developing Children data, Pipeline 1 showed a relatively small number of significant changes, mostly decreases (increases/decreases: 2/17). Pipeline 2 showed a moderate number of significant connectivity increases and only few significant decreases (increases/decreases: 55/5). Pipelines 3 & 4 both showed many significant connectivity increases (increases: 230 and 218) with an absence of significant connectivity decreases.
- In Temporal Lobe Surgery data, Pipeline 1 showed a large number of significant connectivity increases across both hemispheres, along with significant connectivity decreases primarily ipsilaterally (increases/decreases: 325/84). Pipeline 2 showed a comparable number of significant connectivity changes (increases/decreases: 207/254), although the greatest effect sizes were observed for decreases close to resection. Pipelines 3 and 4 primarily showed a high number significant connectivity decreases ipsilateral and proximal to resection, along with several significant increases (increases/decreases: 47/547 [symmetric optimisation] & 38/617 [differential optimisation]); effect sizes were largest in magnitude proximal to the resection.

**Fig. 7.**
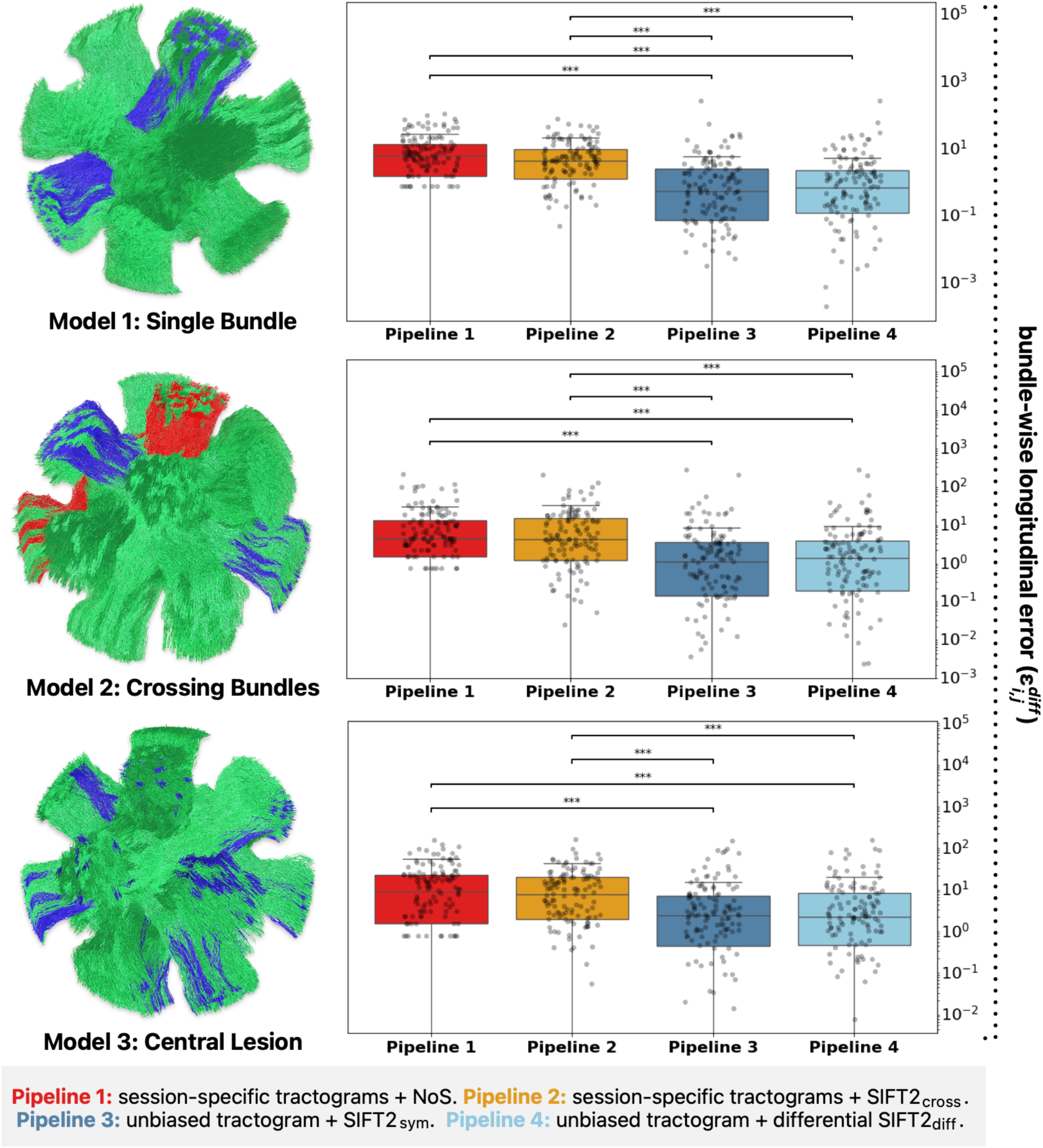
Error in longitudinal structural connectivity quantification. Shown are the pipeline-wise errors in estimated longitudinal fibre count changes across three *in silico* models, with each dot representing a single bundle. Asterisks indicate statistical significance between pipeline pairs (p<10^-4^); statistical testing was performed through the Kruskal-Wallis test with Dunn post-hoc tests, corrected for multiple comparisons. Abbreviations: cross = cross-sectional, diff = differential, SIFT = Spherical-deconvolution Informed Filtering of Tractograms, sym = symmetric.

**Fig 8.**
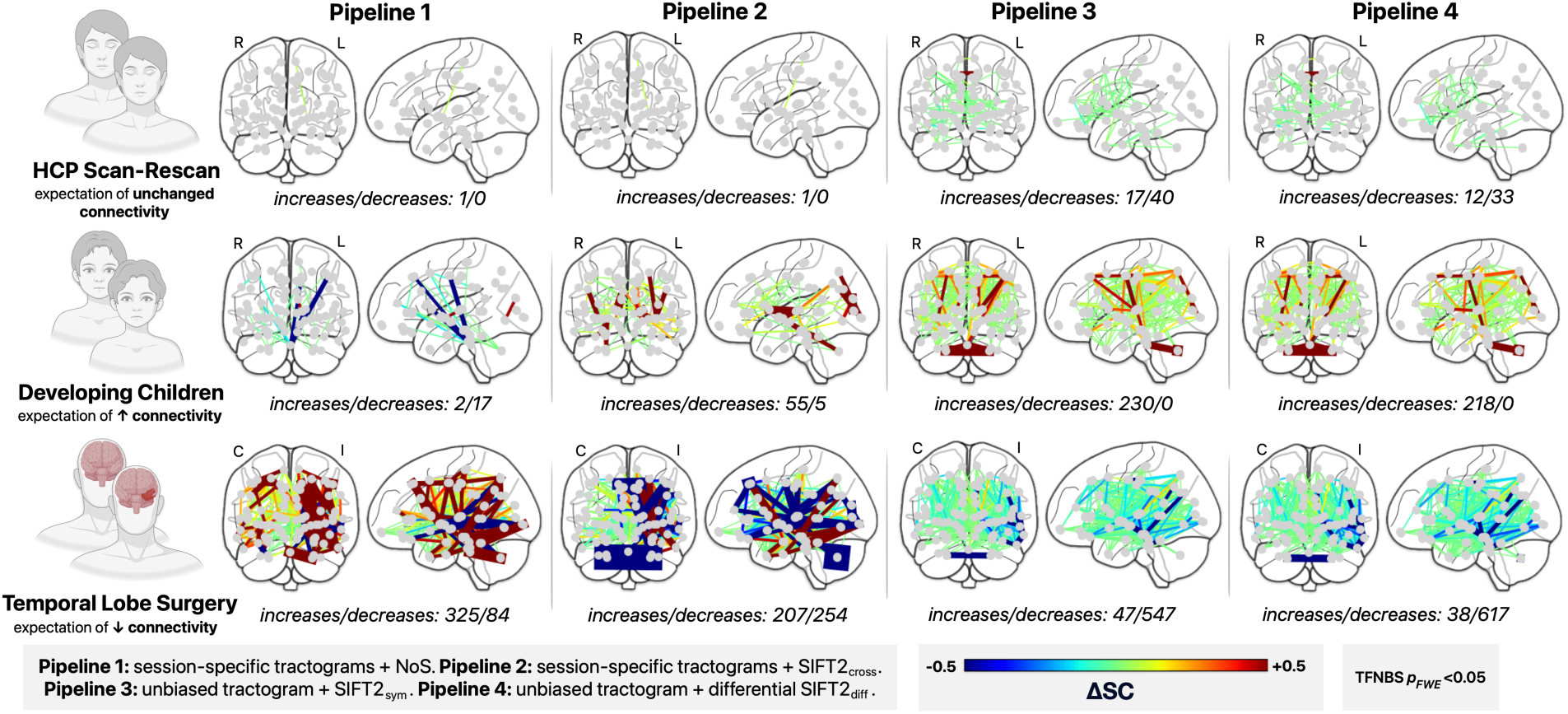
Statistically significant edge-wise FBC changes for *in vivo* cohorts. Shown is the mean change in edge-wise FBC across each cohort and reconstruction pipeline for edges with statistically significant changes identified via TFNBS. Abbreviations: cross = cross-sectional, diff = differential, FWE = Family-Wise Error, HCP = Human Connectome Project, I/C = ipsilateral/contralateral, L/R = left/right, NoS = number of streamlines, SC = structural connectivity, TFNBS = Threshold-Free Network-Based Statistic, SIFT = Spherical-deconvolution Informed Filtering of Tractograms, sym = symmetric.

Altogether, symmetric and differential unbiased reconstruction yielded more extensive findings of statistically significant connectivity changes than cross-sectional pipelines despite using the same statistical framework, indicating greater sensitivity. The identified significant changes were also more concordant with a priori expectations, highlighting improved biological specificity.

## 4 Discussion

In this article, we present a novel quantitative streamline tractography framework tailored to the robust longitudinal quantification of structural connectivity. Unlike conventional pipelines that cross-sectionally run tractography for each session, the proposed framework defines a single quantitative streamline tractogram per individual in an unbiased within-subject template and optimises their ascribed weights to capture connectivity changes over time. We evaluated the framework in the context of longitudinal structural connectome analysis, comparing it against cross-sectional pipelines with and without density optimisation. In *in silico* models, the framework showed significantly lower errors relative to the ground truth. In human *in vivo* data, it yielded statistical outcomes both more extensive and more commensurate with biological expectation. Overall, these findings demonstrate that the proposed method drastically reduces methodological imprecisions when quantifying longitudinal changes in structural connectivity.

### 4.1 The promise and challenges of longitudinal connectivity analysis

Longitudinal studies provide an exciting opportunity to investigate the human brain, offering distinct advantages over traditional case-control studies. Leveraging the consistency in gross anatomy over time, longitudinal study designs have enhanced sensitivity to subtle biological effects that may otherwise be obscured by inter-individual variability. Moreover, exploiting the temporal order of measurements provides a framework for improved causal inference, e.g. observing changes immediately after an intervention can allow more confident attribution of these effects to the intervention.

Despite these conceptual advantages, longitudinal analysis of brain structural connectivity is currently hampered by significant methodological variance(Smith et al., 2015b). Most studies published to date report findings with directionality opposite of what would be expected biologically, e.g. connectivity increases in aging adults(Coelho et al., 2021; Deschwanden et al., 2025), in Parkinson patients(Bergamino et al., 2023), and following surgical resection (Jeong et al., 2016; Ji et al., 2015; Larivière et al., 2024), as well as connectivity decreases in developing adolescents(Baker et al., 2015). These findings not only contradict biological expectation, but also dMRI analyses of similar cohorts with different methods (e.g. Fixel-Based Analysis)(Genc et al., 2018; Han et al., 2023; McDonald et al., 2010; Rau et al., 2019), inculpating technical challenges specific to longitudinal connectivity analysis.

While a comprehensive discussion of the myriad reconstruction biases that can produce spurious connectivity changes is beyond the scope of this article (for a review, see Jeurissen et. al(Jeurissen et al., 2019)), the following sections discuss key challenges in the context of the SIFT framework, and how they can be overcome to robustly estimate changes in Fibre Bundle Capacity (FBC).

### 4.2 Robust attribution of effects to white matter pathways

The application of a tractogram optimisation method such as SIFT2 seeks to correct for reconstruction biases that manifest as local discordances between tractogram and fibre densities. Where applied cross-sectionally to longitudinal data, these specific discordances will be corrected independently per timepoint; one may therefore optimistically expect that, in the presence of some change in local fibre densities between timepoints, a proportional difference in the weights of those streamlines that traverse the same pathway in each of those two timepoints will manifest.

This is however too optimistic with respect to the precision of tractography. Small changes in the local dMRI signal can cause gross differences in the relative proportions of streamlines following different trajectories. These divergences can be far greater in magnitude than the differences in the local fibre densities that serve as the target for tractogram optimisation methods. Since the quantification of bundle connectivity in the context of such methods is still predicated on streamline trajectories, the application of such methods to each timepoint independently is simply incapable of wholly mitigating these imprecisions.

This insight motivated the development of the presented framework, where an individual’s streamline trajectories remain fixed throughout the analysis, allowing only their ascribed weights to vary. The benefits of this approach were quantitatively measured in dMRI phantoms, where we found unbiased pipelines to result in significantly lower errors compared to cross-sectional methods (see Fig 6). Further investigation revealed these improvements to be primarily driven by a reduction of erroneous attribution of effects to false positive pathways (see Supplementary Fig. 5). The rest of this subsection will present additional explanation from first principles and experimental evidence as to why this is the case.

Firstly, it is important to understand that the classification of streamlines as false positives based only on the image data is non-trivial, given that by construction there exists for *every* streamline evidence throughout those image data for its existence(Maier-Hein et al., 2017; Schilling et al., 2022). This limitation is relevant for tractogram optimisation methods, which when tested under realistic ground truth fibre complexities, only yield moderate improvements in terms of false positive filtering(Sarwar et al., 2023), a limitation that applied to *all* evaluated methods within this category, including SIFT(Smith et al., 2013), SIFT2(Smith et al., 2015a), COMMIT(A. Daducci et al., 2015), COMMIT2(Schiavi et al., 2020) and LiFE(Pestilli et al., 2014).

Given the high prevalence of false positives in tractograms(Maier-Hein et al., 2017), and the inability of tractogram optimisation methods to identify them as such(Sarwar et al., 2023), false positive streamlines will inevitably receive non-zero weights. This in return can trigger complex compensatory mechanisms during global optimisation, resulting in unintended errors across the connectome, a behaviour elsewhere compared to the “butterfly-effect” (Bosticardo et al., 2025; Daducci & Schiavi, 2025). The presence of false positive streamlines therefore significantly hinders the accurate quantification of structural connectivity.

Despite the inability of tractogram optimisation to identify false positive streamlines, we here posit that the deleterious effects of their inevitable presence can be greatly diminished where one seeks to quantify connectivity *differences*. Where biological longitudinal effects manifest in a subset of tracts, differences will manifest along the entire length of those streamlines that recapitulate the full trajectory of an affected tract, but only for parts of the length of those that reconstruct unaffected or non-biological pathways. The presented “symmetric” and “differential” optimisation algorithms will therefore ideally tend toward attributing all of the observed local effects to streamlines that reconstruct affected biological pathways; conversely, attributing a non-zero longitudinal effect to a streamline that follows either an erroneous trajectory with no/partial correspondence to a biological tract, or a tract for which there is no biological effect present, will tend to be deleterious to the model fit.

Some empirical evidence for these claims can be obtained from phantom simulations, as they allow explicit experimental control over affected, unaffected and false positive pathways. To explore this, we re-ran parts of the *in silicio* validation (central lesion model) both in the presence and absence of false positive streamlines (around half of the tractogram), reconstructing the first timepoint through absolute optimisation as well as inter-session differences through differential optimisation.

The outcomes of this experiment corroborate several of the above statements (see Fig. 9). The prospective efficacy of absolute and differential tractogram optimisation methods to remove false positive streamlines based on fibre density information alone is undermined by the observation that the optimised tractograms including false positive streamlines had superior fit to the image data than the optimised tractograms excluding such. However, the *consequences* of the presence of false positives were distinctly different between absolute and differential optimisation: the relative increase in errors across bundle categories were substantially smaller for differential reconstruction (simple misattribution of fibre density differences to *some* false positives) as compared to absolute reconstruction (larger misattribution of absolute fibre density to false positive bundles with concomitant errors in true bundles), supporting the prospect of improved attribution of differences in image data to the originating white matter bundles.

**Fig. 9.**
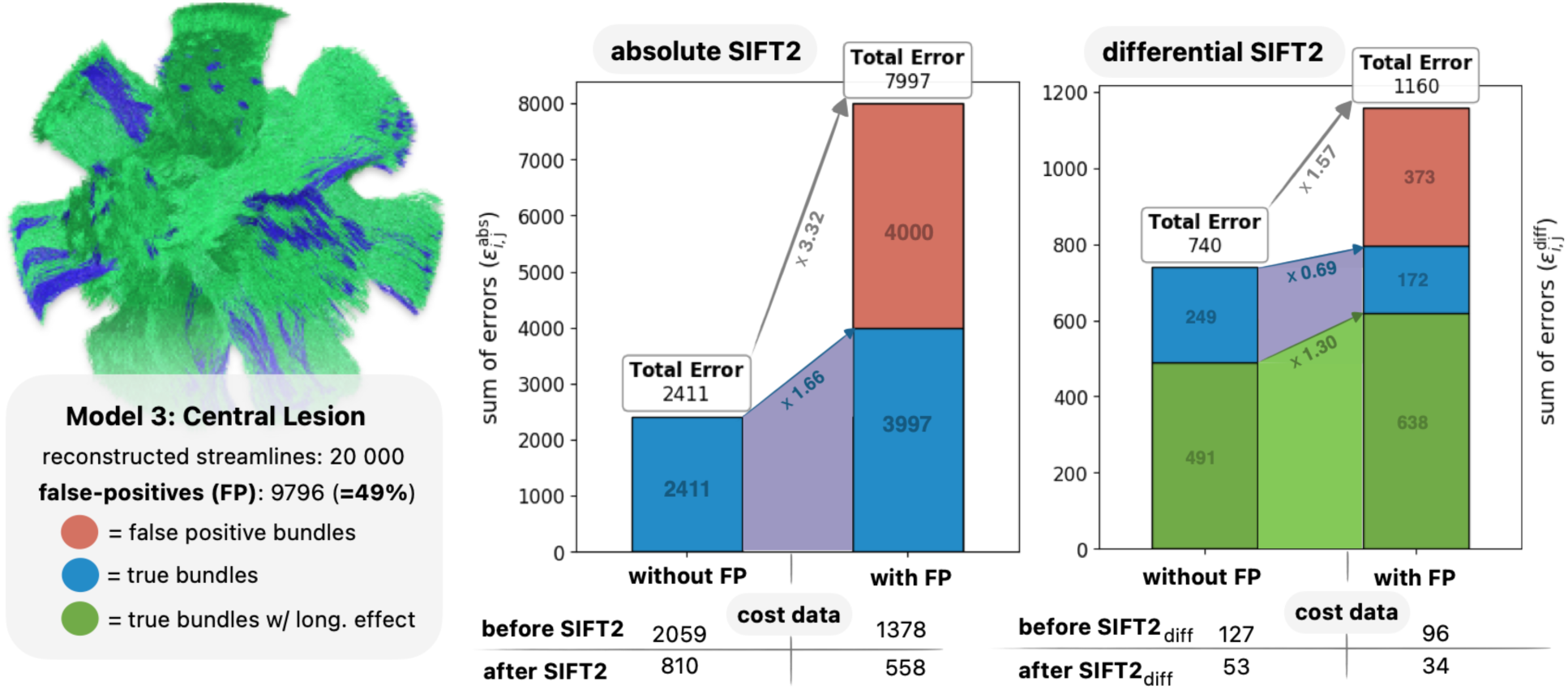
Impact of false positive streamlines on absolute and differential SIFT2 optimisation. Shown are the reconstruction errors within a quantitative tractogram after density optimisation. The left panel shows the magnitude of absolute errors following absolute optimisation of the tractogram at the first timepoint (sum of *ε*^*abs*^, see Supplementary Materials), the right panel shows the magnitude of longitudinal errors (sum of *ε*^*diff*^) following differential optimisation of the unbiased tractogram. The first bar within each panel reports errors if optimisation was performed without any false-positive streamlines, the second bar if optimisation was performed with false-positive streamlines present. The table below the bar plots lists the cost of the data before and after density optimisation. Long = longitudinal.

### 4.3 Validation in human *in vivo* data

While phantom simulations provide a well-controlled environment for numerical testing of connectome pipelines against a known ground truth, they represent only simplified models of biological reality and acquisition complexities. To evaluate the framework in its ultimate target environment, namely human *in vivo* cases (where the ground truth is unknown), we chose three cohorts where there is a strong expectation of what changes to expect biologically. The cohorts were chosen so that they complement each other, together spanning a broad spectrum of expected effects: presence vs. absence of changes, different directionality of changes (increases vs. decreases), and different spatial specificities (widespread vs. localised).

Across all cohorts, unbiased reconstruction pipelines identified larger number of significant edges than their cross-sectional counterparts (see Fig 8), reflecting the notable decrease in edge-wise variance (see Supplementary Fig. 6). For the Developing Children and Temporal Lobe Surgery cohorts, the directionality and spatial distribution of significant connectivity changes obtained from unbiased pipelines conformed well to biological expectation; this was not the case for cross-sectional pipelines, which frequently showed unchanged connectivity where changes would be expected biologically, or changes of biologically non-expected directionality accompanying those that were expected.

To our surprise, all reconstruction pipelines also found statistically significant changes in the HCP Scan-Rescan cohort. While these changes are almost certainly not of biological nature due to the short inter-scan interval (six months), systematic differences may still manifest due to factors beyond our experimental control, such as scanner drift, or different versions of the preprocessing pipeline that were asymmetrically applied to scan and re-scan sessions.

The manifestation of biologically unexpected statistically significant differences – despite stringent correction for multiple comparisons – warranted further investigation. To evaluate the veracity of connectivity changes produced by each pipeline, we generated cohort-average FOD templates, in which we computed the mean fixel-wise change in FDC across subjects following the Fixel-Based Analysis pipeline(Raffelt et al., 2017a). In the same templates, we generated whole-brain tractograms and derived a group-wise parcellation through joint label fusion of subject-specific parcels, allowing the assignment of streamlines to connectome edges. This enabled visual comparison of cohort-average changes in FDC (through fixel colour) and FBC (through streamline colour). Longitudinal connectome pipelines that faithfully reconstruct connectivity changes from local fibre information should produce group-average edge-wise connectivity changes that have at least partial correspondence with group-average changes in local fibre densities. If on the other hand pipelines show effects in areas where the underlying image data suggests no changes or changes in the opposite direction, then this would be suggestive of the quantified effect being the result of tractography biases.

The results of this comparison are shown in Fig. 10, yielding the following observations:

- In HCP-Scan Rescan data, we found mostly unchanged FDC, except for moderate changes of opposing sign along opposing borders of the anterior part of the corpus callosum (the decrease partially overlaps the ventricles and hence may be a registration artifact). Pipelines 1 & 2 showed widespread connectivity changes large in magnitude. Pipelines 3 & 4 showed mostly unchanged connectivity, except for an increase in the edge connecting the bilateral anterior rostral frontal gyri, corresponding well to the FDC increases observed in the corpus callosum.
- In Developing Children data, we found reasonably widespread increases in FDC, most pronounced in the bilateral corticospinal tracts, and minimal decreases in FDC. Pipeline 1 & 2 however showed changes that were even more widespread, with equal proportion of connectivity increases and decreases, most notably with pronounced contradictory decreases in the bilateral corticospinal tracts. Pipelines 3 & 4 showed almost exclusively connectivity increases, including strong effects in the bilateral corticospinal tracts.
- In Temporal Lobe Surgery data, we found decreases in FDC particularly ipsilateral & proximal to resection. Pipeline 1 demonstrated connectivity changes large in magnitude, but non-specific in both sign and hemisphere. Pipeline 2 showed longitudinal changes that were more specific to the ipsilateral hemisphere, but still with a contradictory assortment of both increases and decreases. Pipelines 3 & 4 showed almost exclusively connectivity decreases that were localised ipsilateral & proximal to resection.

**Fig. 10.**
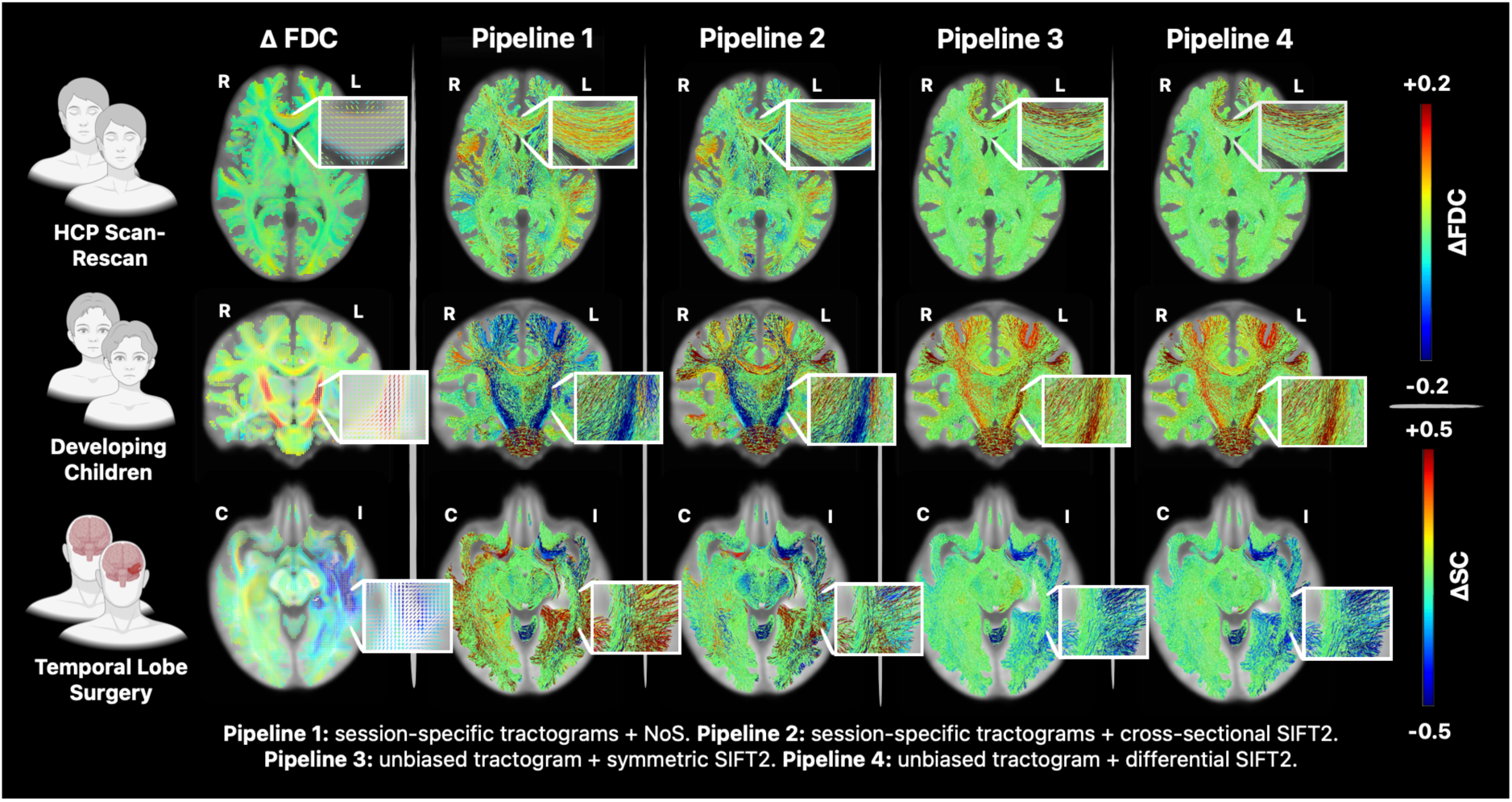
Comparison of edge-wise structural connectivity differences against local fibre density differences. Shown is the mean change in fixel-wise FDC for comparison with the mean change in edge-wise structural connectivity across each cohort and reconstruction pipeline. Abbreviations: FBC = Fibre Bundle Capacity, FDC = Fibre Density and Cross-Section, HCP = Human Connectome Project, I/C = ipsilateral/contralateral, L/R =left/right.

In summary, unbiased pipelines produced connectivity changes that were well supported by the underlying local fibre density estimates, whereas the cross-sectional pipelines reported many connectivity changes of considerable magnitude for which there was no support in the underlying local fibre density estimates. Given that visualisation of the HCP Scan-Rescan image data highlighted a set of fixels with non-zero FDC change concordant with the location of the statistically significant edge reported by the unbiased pipelines, we conclude that this is not an analysis artifact as initially assumed. We further reject the possibility that this observation is due to a systematic bias in the wider image processing pipeline, given that permutation of the session labels in this cohort yielded FDC effects vastly smaller in magnitude and no statistical differences in FBC (data not shown). Together, this is therefore instead suggestive of some systematic bias in image acquisition or pre-processing within this cohort to which the proposed unbiased pipelines are sensitive, the diagnosis of which is beyond the scope of this article.

### 4.4 Importance of adequate connection density normalisation

The quantification of biologically meaningful connectivity differences requires longitudinal measurements to be scaled appropriately for that subtraction operation to be meaningful. Appropriate normalisation of structural connectivity estimates is therefore of imminent relevance for reliable longitudinal comparisons.

The most common strategy for inter-session normalisation is the generation of whole-brain tractograms with a fixed number of streamlines. Absent further manipulation, however, this makes it mathematically impossible for effects to manifest exclusively in one direction: if there is any localised change of one sign in the connectome, there must be a change of the opposite sign elsewhere in the connectome.

This deficiency becomes especially apparent in the case of surgical resection, where there is a gross imbalance in total white matter content between sessions. Despite the clear biological expectation of a degenerative effect, Pipeline 1 (based on streamline count) reported more significant connectivity increases than decreases, many of which were large in magnitude (see Fig. 7). Importantly, these connectivity increases were not supported by cohort-average local fibre density changes (see Fig. 10), indicating these outcomes to be strongly impacted by a failure to adequately normalise across sessions.

The quantification of Fibre Bundle Capacity (FBC) provides appropriate solution to the problem of connection density normalisation(Smith et al., 2022). As part of its model, SIFT2 determines the “proportionality coefficient” *µ*, which serves as a global scaling between tractogram and local fibre densities (see Equation 2). If those local fibre densities are themselves amenable to comparison across sessions (achieved through use of common response functions in the case of constrained spherical deconvolution(Smith et al., 2022)), this global scaling therefore implicitly normalises connectivity estimates so that they become sensitive to biological differences in those local fibre densities between sessions, yet insensitive to the parameters of tractography(Smith et al., 2022).

The relevance of this normalisation is highlighted in a validation study preceding this work: unlike other commonly-used metrics of structural connectivity, longitudinal estimation of FBC appropriately derives the ground truth changes in connectivity regardless of tractography reconstruction configuration(Smith et al., 2025), justifying the adoption of this strategy in the current work.

### 4.5 A cautionary note on brain structural reorganisation

The notion of structural reorganisation has a long-standing history in dMRI research. Following the introduction of the diffusion tensor imaging (DTI) model(Basser et al., 1994), many studies interpreted findings of changes to model anisotropy as a proxy for changes in the “integrity” of underlying connections. While this interpretation was concordant with observations of anisotropy increases following learning tasks(Hofstetter et al., 2013; Scholz et al., 2009; Takeuchi et al., 2010), multiple studies involving neurodegenerative processes also showed localised increases in model anisotropy(Harris et al., 2016; Mole et al., 2016; Yogarajah et al., 2010). Given that such increases were frequently observed alongside biologically expected anisotropy decreases, they were often attributed to compensatory biological mechanisms in response to pathologies, hence reported as signs of “structural reorganisation”. It was only after the introduction of higher-order diffusion models that it became clear that such anisotropy increases could instead result from the presence of complex fibre configurations – subsequently shown to constitute a large fraction of human white matter(Behrens et al., 2007; David et al., 2020; Guo et al., 2020; Jeurissen et al., 2013) – and the inability of the tensor model to resolve such(Jones et al., 2013).

More recently, longitudinal structural connectome studies have similarly reported biologically unexpected changes, which were interpreted as evidence in support of white matter reorganisation(Baker et al., 2015; Bergamino et al., 2023; Coelho et al., 2021; Jeong et al., 2016; Larivière et al., 2024; Mole et al., 2016). Given our understanding of the connection density normalisation problem and the inaccuracies of cross-sectional streamline tractography, there is reasonable concern that such outcomes could also arise as artefacts of inadequate modelling, echoing the prior precedent of erroneous interpretation of DTI-based measures. This is corroborated by findings of the present study, where a naive analysis based on streamline count (i.e. Pipeline 1), involving neither correction of local density biases nor adequate global connection density normalisation, yielded *statistically significant* differences spurious in both sign and location, most prominently in the resection cohort creating the impression of contralateral connectivity increases. Even when appropriately accounting for density biases through tractogram density optimisation, spurious findings remained due to the variability of cross-sectional tractography (i.e. Pipeline 2).

Given this demonstration of both probability and consequence of misleading results from cross-sectional longitudinal connectome reconstruction, the authors urge caution in the interpretation of findings as evidence of brain structural reorganisation—particularly those that contradict current anatomical understanding—and eagerly anticipate future insights arising from the application of more robust analysis methods.

### 4.6 Limitations

The here proposed framework operates on the assumption that the gross white matter architecture remains constant over time, potentially giving it limited sensitivity to the formation of new bundles between timepoints. However, given that white matter bundles are established prenatally(Dubois et al., 2014), we believe this assumption to be well justified for postnatal cohorts.

Generation of unbiased within-subject templates relies on symmetric registration. Failure to accurately align session data within the unbiased image space can have detrimental effects on a wide range of downstream processes executed in this unbiased image space, including but not limited to streamline tractography, computation of fibre density differences, tractogram optimisation and/or the derivation of a brain parcellation. Nonetheless, the here adopted workflow(Avants et al., 2011; Reuter et al., 2012) proved robust across all cases, including those with gross morphological differences between sessions, i.e. surgical resection cases.

The presented differential reconstruction pipeline necessitates mapping of session data to a singular set of fixels for calculation of local fibre density differences. This is presently achieved by segmenting within the template voxel grid both individual session FODs transformed to that space and the FODs of the unbiased template, and subsequently projecting local fibre density values from the former to the latter through the establishment of fixel correspondence(Raffelt et al., 2017b). Small changes to FOD shape could however result in gross differences in fixel segmentation between sessions, which then manifests as fixel fibre density difference disproportionately large relative to any underlying biological change, deleteriously affecting sensitivity to the latter. We here adopted an advanced fixel correspondence strategy to mitigate this effect(Smith & Connelly, 2018), though note that this is not a unique solution(Rafipoor et al., 2025).

Statistical testing of *in vivo* cohorts was performed using the TFNBS framework, which can produce overly extensive results in densely interconnected connectomes(Vinokur et al., 2023), such that not all statistically significant edges are guaranteed to be biologically implicated(Woo et al., 2014). We chose to use TFNBS over stringent mass-univariate comparisons to provide each pipeline with a greater opportunity to detect statistical effects while preserving the spatial specificity of attribution of effects requisite for our assessments. We note that there is no currently accepted best practise for statistical inference in structural connectomes and that this remains an ongoing field of research (Helwegen et al., 2023; Noble & Scheinost, 2020; Serin et al., 2021; Zalesky et al., 2010).

## 5. Conclusion

The here presented quantitative streamline tractography framework enables robust longitudinal quantification of white matter connectivity differences, overcoming key challenges of cross-sectional streamline tractography by optimising a single subject-specific unbiased quantitative tractogram. The substantial performance increases of this framework over conventional cross-sectional pipelines are supported by superior accuracy in *in silico* phantom simulations, as well as by *in vivo* results that were more commensurate with both image data and biological expectation.

## Data and Code Availability

The relevant software to perform symmetrical and differential SIFT2 optimisation is available through the MRtrix3 GitHub repository (https://github.com/orgs/MRtrix3/projects/8/views/1). Data and code to support the findings presented in this study as well as *in silicio* phantoms are available through a separate GitHub repository (https://github.com/ppruc/sift2_unbiased). The repository also contains a comprehensive step-by-step tutorial for users wishing to apply the here presented framework to their own data (https://github.com/ppruc/sift2_unbiased/tutorial). The HCP Scan-Rescan dataset is openly available at https://www.humanconnectome.org/. The Developing Children dataset is openly available at https://openneuro.org/datasets/ds003416 in Brain Imaging Data Structure (BIDS) format with de-identified metadata and defaced images. Access to the Temporal Lobe Surgery dataset is subject to the policies of Vanderbilt University Medical Center and the National Institutes of Health.

## Author contributions

**P.P.:** Conceptualisation; Methodology; Investigation; Formal analysis; Investigation; Visualisation; Writing – original draft; Writing – review & editing.

**R.M.:** Methodology; Supervision; Writing – review & editing.

**D.V.:** Methodology; Supervision; Writing – review & editing.

**K.S.:** Conceptualisation; Resources; Data curation; Writing – review & editing.

**V.M.:** Resources; Data curation; Writing – review & editing.

**D.E.:** Resources; Data curation; Writing – review & editing.

**R.S.:** Conceptualisation; Methodology; Software; Supervision; Project administration; Writing – review & editing.

## Funding

P.P. is supported by the Melbourne International Research Scholarship from the University of Melbourne. R.M. is the recipient of an Australian Research Council Discovery Early Career Researcher Award *(project number: DE240101035).* K.G.S. is supported by National Institutes of Health (NIH) award number K01EB032898. R.E.S. is supported by fellowship funding from the National Imaging Facility *(NIF),* an Australian Government National Collaborative Research Infrastructure Strategy *(NCRIS)* capability. Collection of Temporal Lobe Surgery data supported by National Institutes of Health R01 NS108445 and R01 NS110130. Collection of HCP Scan-Rescan data was funded by the 16 NIH Institutes and Centers that support the NIH Blueprint for Neuroscience Research.

## Ethics statement

Collection of the Developing Children and Temporal Lobe Epilepsy Surgery dataset study was approved by the Vanderbilt University Institutional Review Board, and informed consent was obtained from all participants. HCP Scan-Rescan data were provided by the Human Connectome Project, WU-Minn Consortium (Principal Investigators: David Van Essen and Kamil Ugurbil; <u>1U54MH091657</u>).

## Declaration of Competing Interests

The authors report no competing interests.

## Abbreviations

ACT: Anatomically Constrained Tractography
dMRI: diffusion Magnetic Resonance Imaging
FBC: Fibre Bundle Capacity
FDC: Fibre Density and Cross-section
FOD: Fibre Orientation Distribution
FWE: Family-Wise Error
HCP: Human Connectome Project
IQR: Interquartile Range
ODF: Orientation Distribution Function
SIFT: Spherical-deconvolution Informed Filtering of Tractograms
TFNBS: Threshold-Free Network-Based Statistics
TP: imepoint
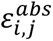: absolute error in structural connectivity
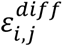: absolute error in longitudinal structural connectivity difference

## Notes

https://github.com/ppruc/sift2_unbiased

### Competing Interest Statement

The authors have declared no competing interest.

### Summary of Updates

Re-upload of the unchanged manuscript PDF to correct formatting issues that appeared after upload to biorxiv, where figures were appended to the end of the document during processing causing scaling issues.

## References

A. Daducci, A. Dal Palù, A. Lemkaddem, & J. -P. Thiran. (2015). COMMIT: Convex Optimization Modeling for Microstructure Informed Tractography. IEEE Transactions on Medical Imaging, 34(1), 246–257. 10.1109/TMI.2014.2352414

Andersson, J. L. R., Skare, S., & Ashburner, J. (2003). How to correct susceptibility distortions in spin-echo echo-planar images: Application to diffusion tensor imaging. NeuroImage, 20(2), 870–888. 10.1016/S1053-8119(03)00336-7

Andersson, J. L. R., & Sotiropoulos, S. N. (2016). An integrated approach to correction for off-resonance effects and subject movement in diffusion MR imaging. NeuroImage, 125, 1063–1078. 10.1016/j.neuroimage.2015.10.019

Avants, B. B., Tustison, N. J., Song, G., Cook, P. A., Klein, A., & Gee, J. C. (2011). A reproducible evaluation of ANTs similarity metric performance in brain image registration. NeuroImage, 54(3), 2033–2044. 10.1016/j.neuroimage.2010.09.025

Baggio, H. C., Abos, A., Segura, B., Campabadal, A., Garcia-Diaz, A., Uribe, C., Compta, Y., Marti, M. J., Valldeoriola, F., & Junque, C. (2018). Statistical inference in brain graphs using threshold-free network-based statistics. Human Brain Mapping, 39(6), 2289–2302. 10.1002/hbm.24007

Baker, S. T. E., Lubman, D. I., Yücel, M., Allen, N. B., Whittle, S., Fulcher, B. D., Zalesky, A., & Fornito, A. (2015). Developmental Changes in Brain Network Hub Connectivity in Late Adolescence. The Journal of Neuroscience, 35(24), 9078. 10.1523/JNEUROSCI.5043-14.2015

Bammer, R., Markl, M., Barnett, A., Acar, B., Alley, M. T., Pelc, N. J., Glover, G. H., & Moseley, M. E. (2003). Analysis and generalized correction of the effect of spatial gradient field distortions in diffusion-weighted imaging. Magnetic Resonance in Medicine, 50(3), 560–569. 10.1002/mrm.10545

Barrios-Martinez, J. V., Fernandes-Cabral, D. T., Abhinav, K., Fernandez-Miranda, J. C., Chang, Y.-F., Suski, V., Yeh, F.-C., & Friedlander, R. M. (2022). Differential tractography as a dynamic imaging biomarker: A methodological pilot study for Huntington’s disease. NeuroImage: Clinical, 35, 103062. 10.1016/j.nicl.2022.103062

Basser, P. J., Mattiello, J., & LeBihan, D. (1994). MR diffusion tensor spectroscopy and imaging. Biophysical Journal, 66(1), 259–267.

Bassett, D. S., & Sporns, O. (2017). Network neuroscience. Nature Neuroscience, 20(3), 353–364. 10.1038/nn.4502

Behrens, T. E. J., Berg, H. J., Jbabdi, S., Rushworth, M. F. S., & Woolrich, M. W. (2007). Probabilistic diffusion tractography with multiple fibre orientations: What can we gain? NeuroImage, 34(1), 144–155. 10.1016/j.neuroimage.2006.09.018

Bergamino, M., Keeling, E. G., Ray, N. J., Macerollo, A., Silverdale, M., & Stokes, A. M. (2023). Structural connectivity and brain network analyses in Parkinson’s disease: A cross-sectional and longitudinal study. Frontiers in Neurology, *Volume 14-*2023. https://www.frontiersin.org/journals/neurology/articles/10.3389/fneur.2023.1137780

Bosticardo, S., Battocchio, M., Schiavi, S., Zalesky, A., Granziera, C., & Daducci, A. (2025). A multi-compartment model for pathological connectomes. Network Neuroscience, 9(4), 1245–1263. 10.1162/NETN.a.30

Cai, Leon Y., Yang, Q., Hansen, C. B., Nath, V., Ramadass, K., Johnson, G. W., Conrad, B. N., Boyd, B. D., Begnoche, J. P., Beason-Held, L. L., Shafer, A. T., Resnick, S. M., Taylor, W. D., Price, G. R., Morgan, V. L., Rogers, B. P., Schilling, K. G., & Landman, B. A. (2021). PreQual: An automated pipeline for integrated preprocessing and quality assurance of diffusion weighted MRI images. Magnetic Resonance in Medicine, 86(1), 456–470. 10.1002/mrm.28678

Cai, Leon Y, Yang, Q., Kanakaraj, P., Nath, V., Newton, A. T., Edmonson, H. A., Luci, J., Conrad, B. N., Price, G. R., & Hansen, C. B. (2021). MASiVar: Multisite, multiscanner, and multisubject acquisitions for studying variability in diffusion weighted MRI. Magnetic Resonance in Medicine, 86(6), 3304–3320.

Coelho, A., Fernandes, H. M., Magalhães, R., Moreira, P. S., Marques, P., Soares, J. M., Amorim, L., Portugal-Nunes, C., Castanho, T., Santos, N. C., & Sousa, N. (2021). Reorganization of brain structural networks in aging: A longitudinal study. Journal of Neuroscience Research, 99(5), 1354–1376. 10.1002/jnr.24795

Cordero-Grande, L., Christiaens, D., Hutter, J., Price, A. N., & Hajnal, J. V. (2019). Complex diffusion-weighted image estimation via matrix recovery under general noise models. NeuroImage, 200, 391–404. 10.1016/j.neuroimage.2019.06.039

Daducci, A., & Schiavi, S. (2025). Quantitative tractography: Joys and sorrows. Brain Structure and Function, 230(6), 88. 10.1007/s00429-025-02939-z

Daducci, A., Schiavi, S., Christiaens, D., Smith, R., & Alexander, D. C. (2025). Chapter 16— Global tractography. In F. Dell’Acqua, M. Descoteaux, & A. Leemans (Eds.), Handbook of Diffusion MR Tractography (pp. 297–314). Academic Press. 10.1016/B978-0-12-818894-1.00014-8

Dale, A. M., Fischl, B., & Sereno, M. I. (1999). Cortical Surface-Based Analysis: I. Segmentation and Surface Reconstruction. NeuroImage, 9(2), 179–194. 10.1006/nimg.1998.0395

David, S., Yousefi Mesri, H., Guo, F., Leemans, A., & De Luca, A. (2020). Diffusion MRI analysis of complex fiber configurations in human brain white matter and cortex.

Deschwanden, P. F., Hotz, I., Mérillat, S., & Jäncke, L. (2025). Functional connectivity-based compensation in the brains of non-demented older adults and the influence of lifestyle: A longitudinal 7-year study. NeuroImage, 308, 121075. 10.1016/j.neuroimage.2025.121075

Desikan, R. S., Ségonne, F., Fischl, B., Quinn, B. T., Dickerson, B. C., Blacker, D., Buckner, R. L., Dale, A. M., Maguire, R. P., Hyman, B. T., Albert, M. S., & Killiany, R. J. (2006). An automated labeling system for subdividing the human cerebral cortex on MRI scans into gyral based regions of interest. NeuroImage, 31(3), 968–980. 10.1016/j.neuroimage.2006.01.021

Dhollander, T., Mito, R., Raffelt, D., & Connelly, A. (2019). Improved white matter response function estimation for 3-tissue constrained spherical deconvolution.

Dhollander, T., Raffelt, D., & Connelly, A. (2016). Unsupervised 3-tissue response function estimation from single-shell or multi-shell diffusion MR data without a co-registered T1 image.

Dubois, J., Dehaene-Lambertz, G., Kulikova, S., Poupon, C., Hüppi, P. S., & Hertz-Pannier, L. (2014). The early development of brain white matter: A review of imaging studies in fetuses, newborns and infants. Secrets of the CNS White Matter, 276, 48–71. 10.1016/j.neuroscience.2013.12.044

Essen, D. C. V., Smith, S. M., Barch, D. M., Behrens, T. E. J., Yacoub, E., & Ugurbil, K. (2013). The WU-Minn Human Connectome Project: An overview. NeuroImage, 80, 62–79. 10.1016/j.neuroimage.2013.05.041

Fischl, B. (2012). FreeSurfer. 20 YEARS OF fMRI, 62(2), 774–781. 10.1016/j.neuroimage.2012.01.021

Garyfallidis, E., Brett, M., Amirbekian, B., Rokem, A., Van Der Walt, S., Descoteaux, M., & Nimmo-Smith, I. (2014). Dipy, a library for the analysis of diffusion MRI data. Frontiers in Neuroinformatics, Volume 8-2014. https://www.frontiersin.org/journals/neuroinformatics/articles/10.3389/fninf.2014.00008

Genc, S., Smith, R. E., Malpas, C. B., Anderson, V., Nicholson, J. M., Efron, D., Sciberras, E., Seal, M. L., & Silk, T. J. (2018). Development of white matter fibre density and morphology over childhood: A longitudinal fixel-based analysis. NeuroImage, 183, 666–676. 10.1016/j.neuroimage.2018.08.043

Glasser, M. F., Sotiropoulos, S. N., Wilson, J. A., Coalson, T. S., Fischl, B., Andersson, J. L., Xu, J., Jbabdi, S., Webster, M., Polimeni, J. R., Van Essen, D. C., & Jenkinson, M. (2013). The minimal preprocessing pipelines for the Human Connectome Project. Mapping the Connectome, 80, 105–124. 10.1016/j.neuroimage.2013.04.127

Guo, F., Leemans, A., Viergever, M. A., Dell’Acqua, F., & De Luca, A. (2020). Generalized Richardson-Lucy (GRL) for analyzing multi-shell diffusion MRI data. NeuroImage, 218, 116948. 10.1016/j.neuroimage.2020.116948

Han, A., Dhollander, T., Sun, Y. L., Chad, J. A., & Chen, J. J. (2023). Fiber-specific age-related differences in the white matter of healthy adults uncovered by fixel-based analysis. Neurobiology of Aging, 130, 22–29. 10.1016/j.neurobiolaging.2023.06.007

Harris, N. G., Verley, D. R., Gutman, B. A., & Sutton, R. L. (2016). Bi-directional changes in fractional anisotropy after experiment TBI: Disorganization and reorganization? NeuroImage, 133, 129–143. 10.1016/j.neuroimage.2016.03.012

Helwegen, K., Libedinsky, I., & van den Heuvel, M. P. (2023). Statistical power in network neuroscience. Trends in Cognitive Sciences, 27(3), 282–301. 10.1016/j.tics.2022.12.011

Hoffmann, M., Salat, D., Reuter, M., & Fischl, B. (2020, August). Longitudinal FreeSurfer with non-linear subject-specific template improves sensitivity to cortical thinning. ISMRM 2020. https://archive.ismrm.org/2020/1050.html

Hofstetter, S., Tavor, I., Moryosef, S. T., & Assaf, Y. (2013). Short-term learning induces white matter plasticity in the fornix. Journal of Neuroscience, 33(31), 12844–12850.

J. G. Sled, A. P. Zijdenbos, & A. C. Evans. (1998). A nonparametric method for automatic correction of intensity nonuniformity in MRI data. IEEE Transactions on Medical Imaging, 17(1), 87–97. 10.1109/42.668698

Jenkinson, M., Beckmann, C. F., Behrens, T. E. J., Woolrich, M. W., & Smith, S. M. (2012). FSL. 20 YEARS OF fMRI, 62(2), 782–790. 10.1016/j.neuroimage.2011.09.015

Jeong, J.-W., Asano, E., Juhász, C., Behen, M. E., & Chugani, H. T. (2016). Postoperative axonal changes in the contralateral hemisphere in children with medically refractory epilepsy: A longitudinal diffusion tensor imaging connectome analysis. Human Brain Mapping, 37(11), 3946–3956. 10.1002/hbm.23287

Jeurissen, B., Descoteaux, M., Mori, S., & Leemans, A. (2019). Diffusion MRI fiber tractography of the brain. NMR in Biomedicine, 32(4), e3785. 10.1002/nbm.3785

Jeurissen, B., Leemans, A., Tournier, J.-D., Jones, D. K., & Sijbers, J. (2013). Investigating the prevalence of complex fiber configurations in white matter tissue with diffusion magnetic resonance imaging. Human Brain Mapping, 34(11), 2747–2766. 10.1002/hbm.22099

Ji, G.-J., Zhang, Z., Xu, Q., Wei, W., Wang, J., Wang, Z., Yang, F., Sun, K., Jiao, Q., Liao, W., & Lu, G. (2015). Connectome Reorganization Associated With Surgical Outcome in Temporal Lobe Epilepsy. Medicine, 94(40). https://journals.lww.com/md-journal/fulltext/2015/10010/connectome_reorganization_associated_with_surgical.41.aspx

Jones, D. K., Knösche, T. R., & Turner, R. (2013). White matter integrity, fiber count, and other fallacies: The do’s and don’ts of diffusion MRI. NeuroImage, 73, 239–254. 10.1016/j.neuroimage.2012.06.081

Jovicich, J., Czanner, S., Greve, D., Haley, E., van der Kouwe, A., Gollub, R., Kennedy, D., Schmitt, F., Brown, G., MacFall, J., Fischl, B., & Dale, A. (2006). Reliability in multi-site structural MRI studies: Effects of gradient non-linearity correction on phantom and human data. NeuroImage, 30(2), 436–443. 10.1016/j.neuroimage.2005.09.046

Larivière, S., Park, B., Royer, J., DeKraker, J., Ngo, A., Sahlas, E., Chen, J., Rodríguez-Cruces, R., Weng, Y., Frauscher, B., Liu, R., Wang, Z., Shafiei, G., Mišić, B., Bernasconi, A., Bernasconi, N., Fox, M. D., Zhang, Z., & Bernhardt, B. C. (2024). Connectome reorganization associated with temporal lobe pathology and its surgical resection. Brain, 147(7), 2483–2495. 10.1093/brain/awae141

Maier-Hein, K. H., Neher, P. F., Houde, J.-C., Côté, M.-A., Garyfallidis, E., Zhong, J., Chamberland, M., Yeh, F.-C., Lin, Y.-C., Ji, Q., Reddick, W. E., Glass, J. O., Chen, D. Q., Feng, Y., Gao, C., Wu, Y., Ma, J., He, R., Li, Q., … Descoteaux, M. (2017). The challenge of mapping the human connectome based on diffusion tractography. Nature Communications, 8(1), 1349. 10.1038/s41467-017-01285-x

McDonald, C. R., Hagler, D. J., Girard, H. M., Pung, C., Ahmadi, M. E., Holland, D., Patel, R. H., Barba, D., Tecoma, E. S., Iragui, V. J., Halgren, E., & Dale, A. M. (2010). Changes in fiber tract integrity and visual fields after anterior temporal lobectomy. Neurology, 75(18), 1631–1638. 10.1212/WNL.0b013e3181fb44db

Mole, J. P., Subramanian, L., Bracht, T., Morris, H., Metzler-Baddeley, C., & Linden, D. E. J. (2016). Increased fractional anisotropy in the motor tracts of Parkinson’s disease suggests compensatory neuroplasticity or selective neurodegeneration. European Radiology, 26(10), 3327–3335. 10.1007/s00330-015-4178-1

Nath, V., Schilling, K. G., Parvathaneni, P., Huo, Y., Blaber, J. A., Hainline, A. E., Barakovic, M., Romascano, D., Rafael-Patino, J., Frigo, M., Girard, G., Thiran, J.-P., Daducci, A., Rowe, M., Rodrigues, P., Prčkovska, V., Aydogan, D. B., Sun, W., Shi, Y., … Landman, B. A. (2020). Tractography reproducibility challenge with empirical data (TraCED): The 2017 ISMRM diffusion study group challenge. Journal of Magnetic Resonance Imaging, 51(1), 234–249. 10.1002/jmri.26794

Neher, P. F., Laun, F. B., Stieltjes, B., & Maier-Hein, K. H. (2014). Fiberfox: Facilitating the creation of realistic white matter software phantoms. Magnetic Resonance in Medicine, 72(5), 1460–1470. 10.1002/mrm.25045

Noble, S., & Scheinost, D. (2020). The Constrained Network-Based Statistic: A New Level of Inference for Neuroimaging. In A. L. Martel, P. Abolmaesumi, D. Stoyanov, D. Mateus, M. A. Zuluaga, S. K. Zhou, D. Racoceanu, & L. Joskowicz (Eds.), Medical Image Computing and Computer Assisted Intervention – MICCAI 2020 (pp. 458– 468). Springer International Publishing.

Pestilli, F., Yeatman, J. D., Rokem, A., Kay, K. N., & Wandell, B. A. (2014). Evaluation and statistical inference for human connectomes. Nature Methods, 11(10), 1058–1063. 10.1038/nmeth.3098

Pruckner, P., Mito, R., Vaughan, D., Jackson, G., Fischmeister, F., Nenning, K.-H., Berger, M., Pataraia, E., Baumgartner, C., Dorfer, C., Rössler, K., Czech, T., Kasprian, G., Bonelli, S., & Smith, R. (2025). Surgical white matter disruption leads to downstream atrophy in the non-resected human brain. *Brain*, awaf344. 10.1093/brain/awaf344

Rafael-Patino, J., Girard, G., Truffet, R., Pizzolato, M., Caruyer, E., & Thiran, J.-P. (2021). The diffusion-simulated connectivity (DiSCo) dataset. Data in Brief, 38, 107429. 10.1016/j.dib.2021.107429

Raffelt, D. A., Tournier, J.-D., Smith, R. E., Vaughan, D. N., Jackson, G., Ridgway, G. R., & Connelly, A. (2017a). Investigating white matter fibre density and morphology using fixel-based analysis. NeuroImage, 144, 58–73. 10.1016/j.neuroimage.2016.09.029

Raffelt, D. A., Tournier, J.-D., Smith, R. E., Vaughan, D. N., Jackson, G., Ridgway, G. R., & Connelly, A. (2017b). Investigating white matter fibre density and morphology using fixel-based analysis. NeuroImage, 144, 58–73. 10.1016/j.neuroimage.2016.09.029

Raffelt, D., Tournier, J-Donald, Crozier, S., Connelly, A., & Salvado, O. (2012). Reorientation of fiber orientation distributions using apodized point spread functions. Magnetic Resonance in Medicine, 67(3), 844–855. 10.1002/mrm.23058

Raffelt, D., Tournier, J.-Donald, Rose, S., Ridgway, G. R., Henderson, R., Crozier, S., Salvado, O., & Connelly, A. (2012). Apparent Fibre Density: A novel measure for the analysis of diffusion-weighted magnetic resonance images. NeuroImage, 59(4), 3976– 3994. 10.1016/j.neuroimage.2011.10.045

Rafipoor, H., Lange, F. J., Arthofer, C., Cottaar, M., & Jbabdi, S. (2025). Hierarchical modelling of crossing fibres in the white matter. Imaging Neuroscience, 3, imag_a_00436. 10.1162/imag_a_00436

Rau, Y.-A., Wang, S.-M., Tournier, J.-D., Lin, S.-H., Lu, C.-S., Weng, Y.-H., Chen, Y.-L., Ng, S.-H., Yu, S.-W., Wu, Y.-M., Tsai, C.-C., & Wang, J.-J. (2019). A longitudinal fixel-based analysis of white matter alterations in patients with Parkinson’s disease. NeuroImage: Clinical, 24, 102098. 10.1016/j.nicl.2019.102098

Reuter, M., & Fischl, B. (2011). Avoiding asymmetry-induced bias in longitudinal image processing. NeuroImage, 57(1), 19–21. 10.1016/j.neuroimage.2011.02.076

Reuter, M., Schmansky, N. J., Rosas, H. D., & Fischl, B. (2012). Within-subject template estimation for unbiased longitudinal image analysis. NeuroImage, 61(4), 1402–1418. 10.1016/j.neuroimage.2012.02.084

Sarica, A., Gramigna, V., Arcuri, F., Crasà, M., Calomino, C., Nisticò, R., Bianco, M. G., Quattrone, Andrea, & Quattrone, Aldo. (2025). Differential tractography identifies a distinct pattern of white matter alterations in essential tremor with or without resting tremor. NeuroImage: Clinical, 45, 103734. 10.1016/j.nicl.2025.103734

Sarwar, T., Ramamohanarao, K., Daducci, A., Schiavi, S., Smith, R. E., & Zalesky, A. (2023). Evaluation of tractogram filtering methods using human-like connectome phantoms. NeuroImage, 281, 120376. 10.1016/j.neuroimage.2023.120376

Schiavi, S., Ocampo-Pineda, M., Barakovic, M., Petit, L., Descoteaux, M., Thiran, J.-P., & Daducci, A. (2020). A new method for accurate in vivo mapping of human brain connections using microstructural and anatomical information. Science Advances, 6(31), eaba8245. 10.1126/sciadv.aba8245

Schilling, K. G., Blaber, J., Huo, Y., Newton, A., Hansen, C., Nath, V., Shafer, A. T., Williams, O., Resnick, S. M., Rogers, B., Anderson, A. W., & Landman, B. A. (2019). Synthesized b0 for diffusion distortion correction (Synb0-DisCo). Artificial Intelligence in MRI, 64, 62–70. 10.1016/j.mri.2019.05.008

Schilling, K. G., Tax, C. M. W., Rheault, F., Landman, B. A., Anderson, A. W., Descoteaux, M., & Petit, L. (2022). Prevalence of white matter pathways coming into a single white matter voxel orientation: The bottleneck issue in tractography. Human Brain Mapping, 43(4), 1196–1213. 10.1002/hbm.25697

Scholz, J., Klein, M. C., Behrens, T. E. J., & Johansen-Berg, H. (2009). Training induces changes in white-matter architecture. Nature Neuroscience, 12(11), 1370–1371. 10.1038/nn.2412

Serin, E., Zalesky, A., Matory, A., Walter, H., & Kruschwitz, J. D. (2021). NBS-Predict: A prediction-based extension of the network-based statistic. NeuroImage, 244, 118625. 10.1016/j.neuroimage.2021.118625

Smith, R., & Connelly, A. (2018). Mitigating the effects of imperfect fixel correspondence in Fixel-Based Analysis.

Smith, R. E., Tournier, J.-D., Calamante, F., & Connelly, A. (2012). Anatomically-constrained tractography: Improved diffusion MRI streamlines tractography through effective use of anatomical information. NeuroImage, 62(3), 1924–1938. 10.1016/j.neuroimage.2012.06.005

Smith, R. E., Tournier, J.-D., Calamante, F., & Connelly, A. (2013). SIFT: Spherical-deconvolution informed filtering of tractograms. NeuroImage, 67, 298–312. 10.1016/j.neuroimage.2012.11.049

Smith, R. E., Tournier, J.-D., Calamante, F., & Connelly, A. (2015a). SIFT2: Enabling dense quantitative assessment of brain white matter connectivity using streamlines tractography. NeuroImage, 119, 338–351. 10.1016/j.neuroimage.2015.06.092

Smith, R. E., Tournier, J.-D., Calamante, F., & Connelly, A. (2015b). The effects of SIFT on the reproducibility and biological accuracy of the structural connectome. NeuroImage, 104, 253–265. 10.1016/j.neuroimage.2014.10.004

Smith, R., Pruckner, P., & Abbott, D. (2025). Fibre Bundle Capacity addresses structural connectivity biases in surgical resection cohorts. 10.31219/osf.io/evsb2_v1

Smith, R., Raffelt, D., Tournierc, J.-D., & Connelly, A. (2022). Quantitative streamlines tractography: Methods and inter-subject normalisation. Aperture Neuro, 1–25. 10.52294/ApertureNeuro.2022.2.NEOD9565

Smith, R., Skoch, A., Bajada, C., Caspers, S., & Connelly, A. (2020). Hybrid Surface-Volume Segmentation for improved Anatomically-Constrained Tractography.

Takeuchi, H., Sekiguchi, A., Taki, Y., Yokoyama, S., Yomogida, Y., Komuro, N., Yamanouchi, T., Suzuki, S., & Kawashima, R. (2010). Training of working memory impacts structural connectivity. Journal of Neuroscience, 30(9), 3297–3303.

Tournier, J.-D., Calamante, F., & Connelly, A. (2010). Improved probabilistic streamlines tractography by 2nd order integration over fibre orientation distributions. Proc. Intl. Soc. Mag. Reson. Med. (ISMRM), 18.

Tournier, J.-D., Smith, R., Raffelt, D., Tabbara, R., Dhollander, T., Pietsch, M., Christiaens, D., Jeurissen, B., Yeh, C.-H., & Connelly, A. (2019). MRtrix3: A fast, flexible and open software framework for medical image processing and visualisation. NeuroImage, 202, 116137. 10.1016/j.neuroimage.2019.116137

Veraart, J., Novikov, D. S., Christiaens, D., Ades-aron, B., Sijbers, J., & Fieremans, E. (2016). Denoising of diffusion MRI using random matrix theory. NeuroImage, 142, 394–406. 10.1016/j.neuroimage.2016.08.016

Vinokur, L., Smith, R., Dhollander, T., Vaughan, D., Jackson, G., & Connelly, A. (2023). Parameter Sensitivity of Network-Based Statistical Inference. 10.21203/rs.3.rs-3081615/v1

Vinokur, L., Zalesky, A., Raffelt, D., Smith, R., & Connelly, A. (2015a). A Novel Threshold -Free Network-Based Statistics Method: Demonstration using Simulated Pathology.

Vinokur, L., Zalesky, A., Raffelt, D., Smith, R., & Connelly, A. (2015b). A NOVEL THRESHOLD-FREE NETWORK-BASED STATISTICAL METHOD: DEMONSTRATION AND PARAMETER OPTIMISATION USING IN VIVO SIMULATED PATHOLOGY.

Winkler, A. M., Ridgway, G. R., Webster, M. A., Smith, S. M., & Nichols, T. E. (2014). Permutation inference for the general linear model. NeuroImage, 92, 381–397. 10.1016/j.neuroimage.2014.01.060

Woo, C.-W., Krishnan, A., & Wager, T. D. (2014). Cluster-extent based thresholding in fMRI analyses: Pitfalls and recommendations. NeuroImage, 91, 412–419. 10.1016/j.neuroimage.2013.12.058

Yeh, F.-C., Zaydan, I. M., Suski, V. R., Lacomis, D., Richardson, R. M., Maroon, J. C., & Barrios-Martinez, J. (2019). Differential tractography as a track-based biomarker for neuronal injury. NeuroImage, 202, 116131. 10.1016/j.neuroimage.2019.116131

Yendiki, A., Reuter, M., Wilkens, P., Rosas, H. D., & Fischl, B. (2016). Joint reconstruction of white-matter pathways from longitudinal diffusion MRI data with anatomical priors. NeuroImage, 127, 277–286. 10.1016/j.neuroimage.2015.12.003

Yogarajah, M., Focke, N. K., Bonelli, S. B., Thompson, P., Vollmar, C., McEvoy, A. W., Alexander, D. C., Symms, M. R., Koepp, M. J., & Duncan, J. S. (2010). The structural plasticity of white matter networks following anterior temporal lobe resection. Brain, 133(8), 2348–2364. 10.1093/brain/awq175

Zalesky, A., Fornito, A., & Bullmore, E. T. (2010). Network-based statistic: Identifying differences in brain networks. NeuroImage, 53(4), 1197–1207. 10.1016/j.neuroimage.2010.06.041

