## Supplementary Materials for "Longitudinal quantitative streamline tractography: robust estimation of white matter connectivity differences"

### Supplementary Methods

#### Regularisation functions and bases

The regularisation terms within the original absolute ($R^{abs}$) and the novel differential implementation ($R^{diff}$) mitigate the derivation of optimised solutions where the underlying signal is explained by a small number of streamlines with extreme values, as this incurs quantisation artifacts comparable to having reconstructed too few streamlines. While three suitable regularisations for the original absolute SIFT2 implementation were previously presented in conjunction with that method(Smith et al., 2015), they are here refactored and generalised to facilitate both expansion of options and extensions tailored for differential optimisation.

A chosen configuration of regularisation, whether intended for absolute or differential mode optimisation, consists of two attributes:

1. The regularisation "basis": This is the set of elements upon which regularisation is applied. These bases are mutually common between absolute and differential optimisations.
2. The "functional form" of the regularisation: This is an algebraic expression that translates per-streamline values being optimised to the corresponding cost function penalty. These are tailored to absolute and differential optimisations individually.

The upcoming sections describe the relevant concepts and implementations of regularisation bases and functions.

#### Regularisation bases

Key to an understanding of the regularisation bases is the notion of a "reference" value, wherein regularisation is applied to streamlines based on the magnitude of the deviation of their ascribed value from that reference.

- *Streamline*-wise reference: The regularisation of each streamline is independent of all others. This is equivalent mathematically to a naïve reference value (this will be shown explicitly in each instance).
- *Fixel*-wise reference: For each fixel, the corresponding mean value across those streamlines traversing it is computed; the value ascribed to each streamline is then discouraged from deviating from the mean values within the set of fixels traversed by that streamline.
- *Group*-wise reference: Streamlines are pre-allocated into groups; this could be based on e.g. geometric proximity of streamlines, or common attribution to a particular connectome edge. The mean per-streamline value is computed for each group individually, and the values ascribed to each streamline is then discouraged from deviating from the corresponding mean.

#### Regularisation functions

Given a chosen regularisation basis, regularisation functions compute the penalty for streamline coefficients (absolute) or delta-coefficients (differential) as they deviate from a reference value. The functions are tailored to each mode of optimisation and vary in how strictly they incentivise the algorithm to produce solutions with the desired characteristics.

##### Absolute optimisation

For regularisation of absolute optimisation, three regularisation functions were presented in the original SIFT2 paper(Smith et al., 2015), which are here termed "coefficient", "factor" and "gamma" function. While the coefficient function imposes a penalty directly on the exponential coefficient ($F_s$) as it deviates from a reference point, the factor function imposes a heavier penalty on deviating factors ($e^{F_s}$). The gamma function combines these two functions, choosing to either regularise using the coefficient or factor function, dependent on the value of the exponential coefficient.

#### Differential optimisation

For regularisation of differential optimisation, upcoming sections will describe two regularisation functions, which we term "delta" and "dual inverse barrier" functions. Both behave in a similar manner, penalising delta coefficients ($\Delta F_s$) as they deviate from zero; while the delta function imposes a soft constraint that penalises deviations, the dual inverse barrier function strictly forces delta coefficients to remain within the target range of (-1,1).

#### Regularisation configurations absolute SIFT2

The following subsections outline different configurations of the regularisation bases and functions for absolute optimisation.

##### Streamline basis + coefficient function

The regularisation cost of the coefficient function applied on a streamline basis is computed as follows:

$$R^{abs}(F_s) = (F_s)^2$$

, where $F_s$ denotes the exponential coefficient of streamline *s*. This formulation imposes a quadratically increasing penalty as coefficients deviate from zero, thereby keeping streamline weights ($e^{F_s}$) close to one (see Supplementary Fig. 1, "coefficient function" + "naïve reference").

##### Streamline basis + factor function

The regularisation cost of the factor function applied on a streamline basis is computed as follows:

$$R^{abs}(F_s) = R(e^{F_s} - 1)^2$$

, where $e^{Fs}$ denotes the streamline weight for streamline *s*. In this case, the regularisation penalty quadratically increases as factors deviate from one (see Supplementary Fig. 1, "factor function" + "naïve reference"). Given that streamline factors are the exponential transform of streamline coefficients, the factor function penalises increases in coefficients more heavily than the coefficient function.

#### Streamline basis + gamma function

The regularisation cost of the gamma function applied on a streamline basis is computed the follows:

$$R^{abs}(F_s) = \begin{cases} (F_s)^2, & if\ F_s \leq 0 \\ (e^{F_s} - 1)^2, & if\ F_s > 0 \end{cases}$$

, where $F_s$ denotes the exponential coefficient of streamline $s$, and $e^{Fs}$ denotes the streamline weight of streamlines $s$. The regularisation cost is computed through either the coefficient or factor function, dependent on the value of the exponential coefficient, imposing heavier penalties for $F_s > 0$ (see Supplementary Fig. 1, "gamma function" + "naïve reference").

#### Fixel/group basis + coefficient function

The regularisation cost of the coefficient function applied on a fixel, or a group basis is computed as follows:

$$R^{abs}(F_s, F_{ref}) = (F_s - F_{ref})^2$$

, where $F_s$ denotes the exponential coefficient of streamline $s$, and $F_{ref}$ the mean exponential coefficient of all streamlines assigned to a reference fixel or group. The formulation imposes a quadratically increasing penalty as streamline coefficients deviate from the local mean (see Supplementary Fig. 1, "coefficient function" + "shifting references").

##### Fixel/group basis + factor function

The regularisation cost of the factor function applied on a fixel, or group basis is computed as follows:

$$R^{abs}(F_s, F_{ref}) = (e^{F_s} - e^{F_{ref}})^2$$

, where $e^{F_s}$ is the streamline weight of streamline *s,* and $e^{F_{ref}}$ is the mean streamline weight of streamlines assigned to a reference fixel or group. Here, the regularisation penalty quadratically increases as factors deviate from the local mean (see Supplementary Fig. 1, "factor function" + "shifting references"). As previously, the factor function penalises increases in streamline coefficients more heavily than the coefficient function.

##### Fixel/group basis + gamma function

The regularisation cost of the *gamma* function applied on a fixel, or group basis is computed as follows:

$$R^{abs}(F_s, F_{ref}) = \begin{cases} (F_s - F_{ref})^2, \text{if } F_s \leq F_{ref} \\ (e^{F_s} - e^{F_{ref}})^2, \text{if } F_s > F_{ref} \end{cases}$$

, where $F_s$ denotes the exponential coefficient assigned to streamline *s*, $e^{Fs}$ denotes the streamline weight of streamline *s,* $F_{ref}$ is the mean exponential coefficient of streamlines assigned to a reference fixel or group, and $e^{F_{ref}}$ is the mean streamline weight of streamlines assigned to either a reference fixel or group. Regularisation cost is computed through either the coefficient or factor function, dependent on the value of the exponential coefficient. Heavier penalties are imposed for $F_s > F_{ref}$ (see Supplementary Fig. 1, "gamma function" + "shifting references").

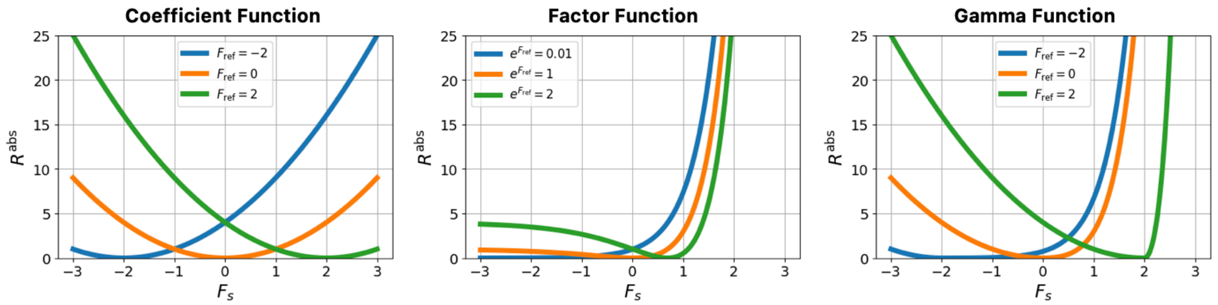


**Supplementary Fig. 1 – Absolute regularisation functions.** Shown are the different functions to compute regularisation costs for absolute optimisation at different reference values. The orange line shows the naïve reference with either $F_{ref} = 0$ or $e^{F_{ref}} = 1$, blue and green lines show how the regularisation behaviour adapts as the target reference shifts from the naïve reference. Symbols: $e^{F_{ref}}$ = mean streamline weight of streamlines assigned to a reference, $F_s$ = streamline coefficient of streamline $s$, $F_{ref}$ = mean streamline coefficient of streamlines assigned to a reference, $R^{abs}$ = absolute regularisation cost.

#### Regularisation differential SIFT2

The following subsections outline different configurations of regularisation bases and functions for differential optimisation.

##### Streamline basis + delta function

The regularisation cost of the delta function applied on a streamline basis is computed as follows:

$$R^{diff}(\Delta F_s) = \Delta F_s^2$$

, where $\Delta F_s$ denotes the delta coefficient associated with streamline $s$, bounded to the range $\Delta F_s \in [-1,1]$. This regularisation imposes a quadratically increasing penalty on delta coefficients as they deviate from zero (see Supplementary Fig. 2, "delta function" + "naïve reference").

##### Streamline basis + dual inverse barrier function

The regularisation cost of the dual inverse barrier applied on a streamline basis is computed as follows:

$$R^{diff}(\Delta F_s) = -1 - \frac{1}{(\Delta F_s - 1)(\Delta F_s + 1)}$$

, where $\Delta F_s$ denotes the delta coefficient associated with streamline $s$, bounded to the range $\Delta F_s \in (-1,1)$. This function penalises values that deviate from zero more heavily, causing the regularisation cost to blow up towards infinity as delta coefficients approximate ±1, strictly keeping them within that range (see Supplementary Fig. 2, "dual inverse barrier function" + "naïve reference").

#### Fixel/group basis + delta function

The regularisation cost of the delta function applied on a fixel/group basis is computed as follows:

$$R^{diff}(\Delta F_s, \Delta F_{ref}) = \begin{cases} (\frac{\Delta F_s - \Delta F_{ref}}{1 + \Delta F_{ref}})^2, & if\ \Delta F_s \leq \Delta F_{ref} \\ (\frac{\Delta F_s - \Delta F_{ref}}{1 - \Delta F_{ref}})^2, & if\ \Delta F_s > \Delta F_{ref} \end{cases}$$

, where $\Delta F_s$ is the delta coefficient of streamline *s*, bounded to the range $\Delta F_s \in [-1,1]$; $\Delta F_{ref}$ is the mean delta coefficient of streamlines assigned to a reference fixel or group. Here, delta coefficients of streamlines are penalised as they deviate from the local mean (see Supplementary Fig. 2, “delta function” + “shifting references”).

#### Fixel/group basis + dual inverse barrier function

The regularisation cost of the dual inverse barrier function applied to on fixel/group basis is computed as follows:

$$R^{diff}(\Delta F_s, \Delta F_{ref}) = \begin{cases} -1 - \frac{1}{(X-1)(X+1)}\ where\ X = \frac{\Delta F_s + 1}{1 + \Delta F_{ref}} - 1, & if\ \Delta F_s \leq \Delta F_{ref} \\ -1 - \frac{1}{(X-1)(X+1)}\ where\ X = \frac{\Delta F_s - 1}{1 - \Delta F_{ref}} + 1, & if\ \Delta F_s > \Delta F_{ref} \end{cases}$$

, where $\Delta F_s$ is the delta coefficient of streamline *s*, bounded to the range $\Delta F_s \in (-1,1)$; $\Delta F_{ref}$ is the mean delta coefficient of streamlines assigned to a reference fixel or group. This regularisation strategy heavily penalises delta coefficients as they deviate from the local mean, while keeping them strictly within the target range (see Supplementary Fig 2, “dual inverse barrier function” + “shifting references”).

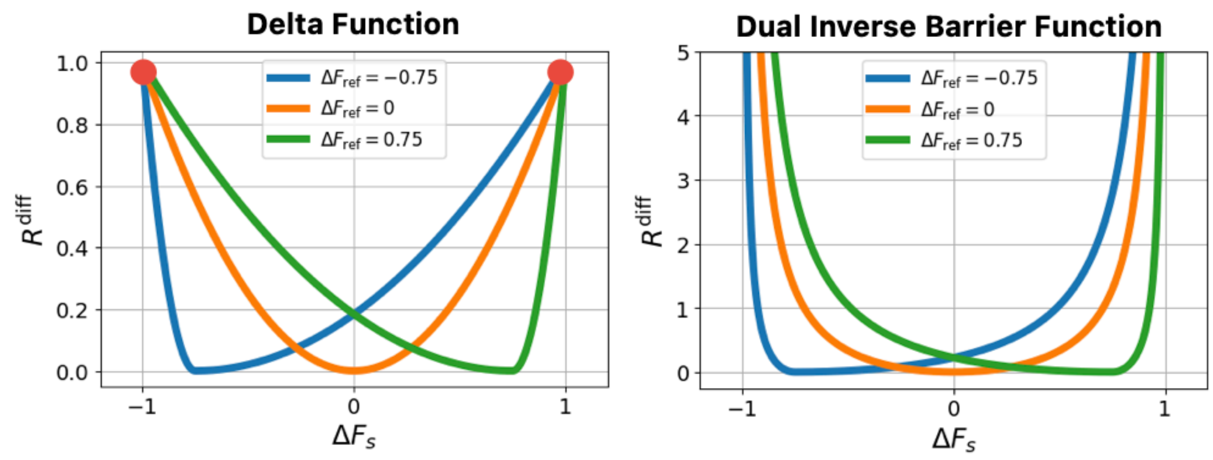


**Supplementary Fig. 2 – Differential regularisation functions.** Shown are the different functions to compute regularisation costs for differential optimisation at different reference values. The orange line shows the naïve reference with $\Delta F_{ref} = 0$, blue and green lines show how the regularisation behaviour adapts if the target reference becomes non-zero. Note that while the delta function is finitely bound within the range [-1,1] (red dots), the dual inverse barrier function asymptotically approaches the bounds (-1,1). Symbols: $\Delta F_s$ = delta coefficient of streamline $s$, $\Delta F_{ref}$ = mean delta coefficient of streamlines assigned to a fixel or group, $R^{diff}$ = regularisation cost.

#### Choosing the appropriate regularisation strength

The strength of regularisation can be modulated through the user-controllable parameter "λ"; to determine an appropriate regularisation strength, a common approach is to perform an L-curve analysis, determining the maximum regularisation strength before which data residuals start to increase. However as shown in this article, a mathematically optimal solution does not necessarily translate to the biologically most accurate solution (see Fig. 9).

This prompted us to perform an extended number of analyses to determine an appropriate regularisation strength for absolute and differential optimisation:

- L-curve analyses, comparing the fit of the solution against the underlying (noisy) local fibre densities.
- Inspection of streamline weight/coefficient distributions, determining the threshold from which on solutions are no longer dominated by only few streamlines with extreme values.
- Assessment of errors in fibre count (change) quantification, comparing the derived solution against the underlying ground truth.

Evaluations were performed in the central lesion model phantom (see Methods, "Validation in *in silico* dMRI phantoms"), and where possible, in an exemplar *in vivo* subject from the Developing Children cohort (see Methods, "Validation in human *in vivo* cohorts"). We assessed the behaviour of all regularisation configurations at different regularisation strengths. For group-wise regularisation, streamlines were grouped based on their assignment to connectome parcels.

##### L-Curve Analyses

L-curve analyses were performed as described in the original SIFT2 publication(Smith et al., 2015). In short, the residual data cost (= the total error across all fixels after optimisation) were plotted against the total regularisation cost (= the total cost of $R^{abs}/R^{diff}$ across all streamlines after optimisation) in log-log space. This allows visual identification of the regularisation strength that best balances accurate data fit with the desired smoothness enforced by regularisation.

The results of this analysis are shown in Supplementary Figure 3 (absolute optimisation) and Supplementary Figure 4 (differential optimisation), and can be summarised as follows:

- The L-curve criterion for absolute SIFT2 optimisation in our central lesion phantom and in vivo subject indicated the ideal regularisation strength to lie at approximately $10^{-3}$ for the different configurations of regularisation bases and functions.
- The L-curve criterion for differential SIFT2 optimisation in our central lesion phantom and in vivo subject indicated the ideal regularisation strength to lie at approximately $10^{-3}$ for configurations including the delta function, and approximately $10^{-4}$ for configurations including the dual inverse barrier function.

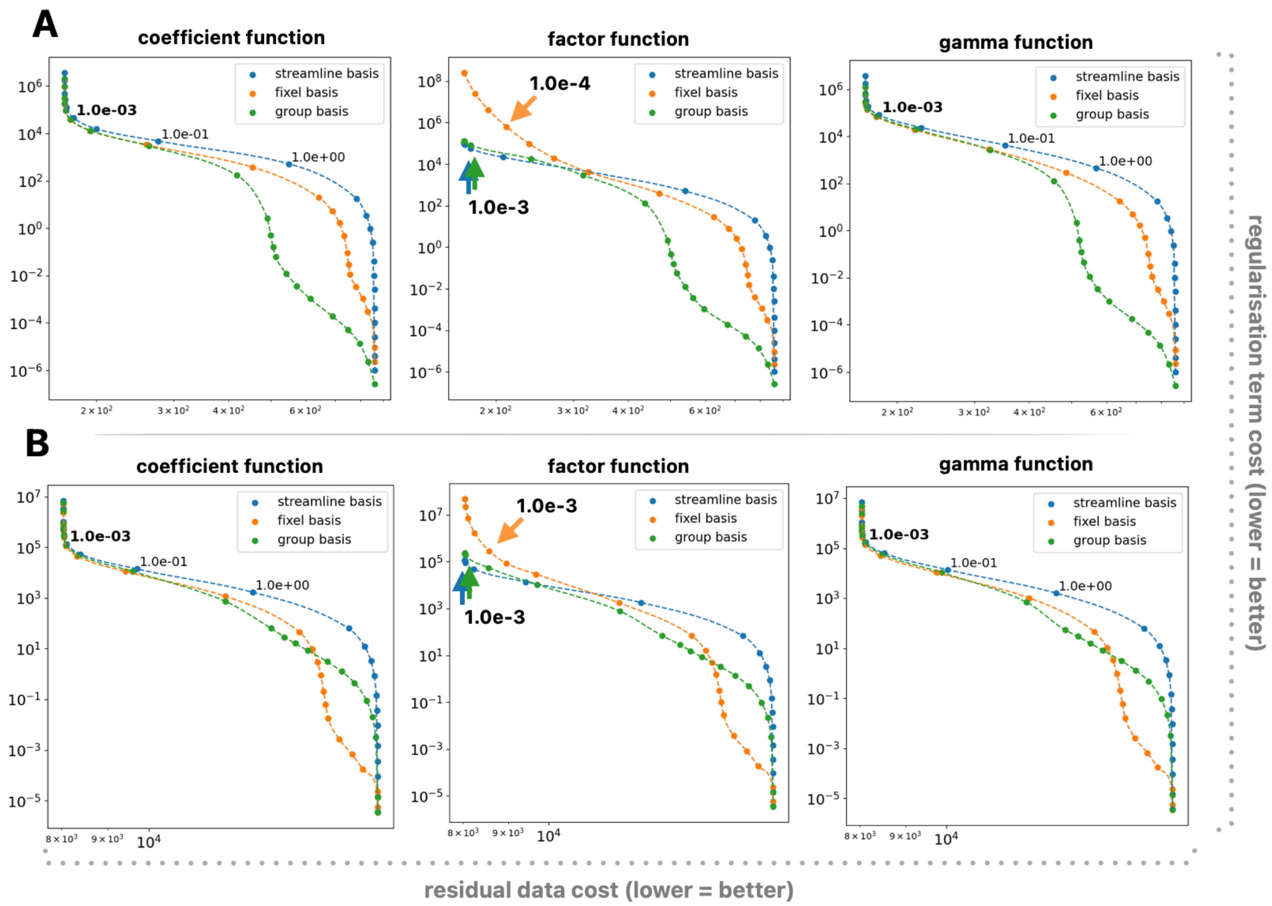


**Supplementary Fig 3 - L-Curves of absolute regularisation functions.** Shown are the L-curves for different absolute regularisation bases and functions, with A) showing L-curves computed on phantom and B) showing L-curves computed on an in vivo subject. Printed in bold is the strongest absolute regularisation strength before data residuals started to increase.

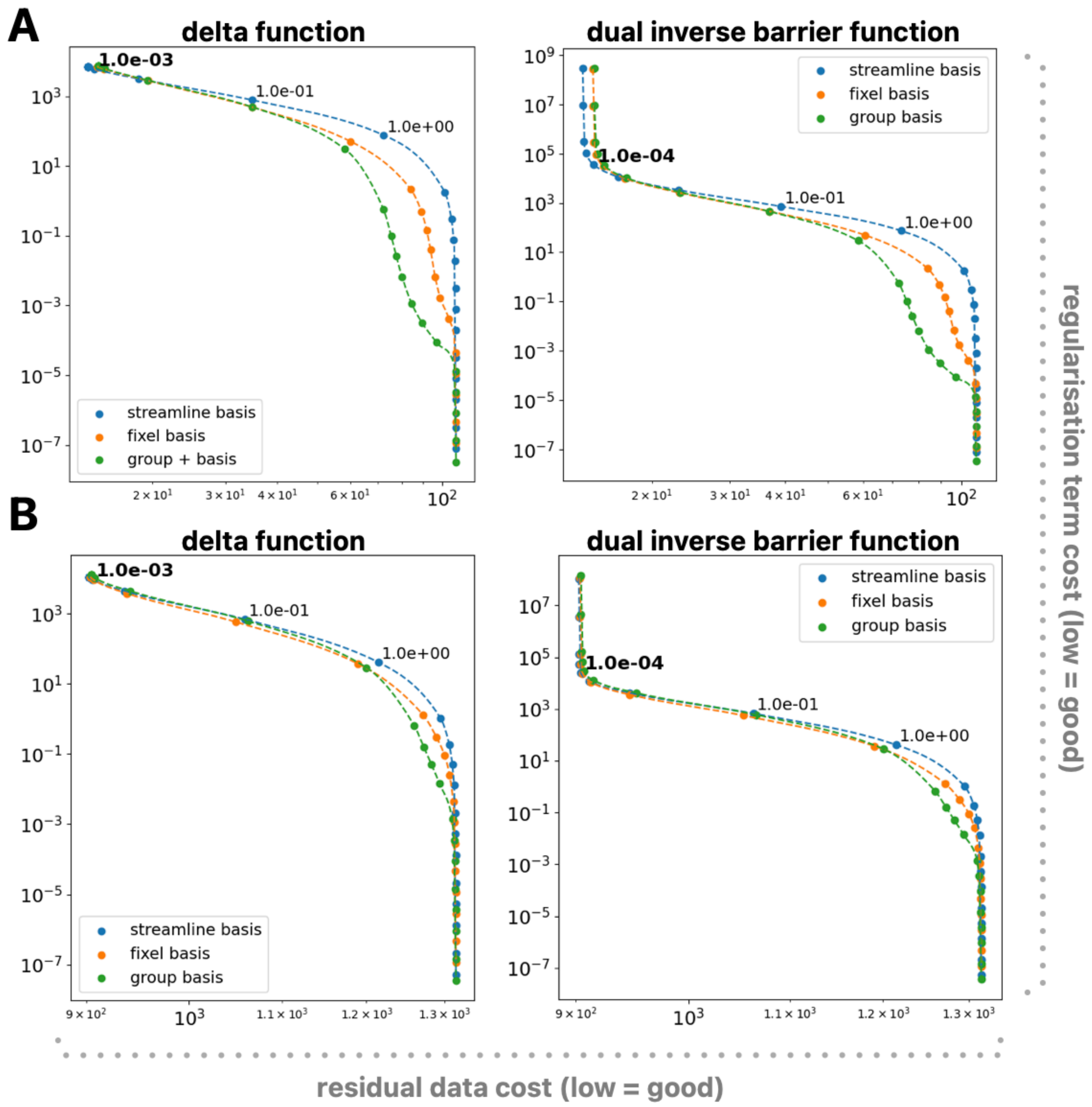


**Supplementary Fig 4 - L-Curves of differential regularisation functions.** Shown are the L-curves for differential regularisation bases and functions, with A) showing L-curves computed on phantom and B) showing L-curves computed on an in vivo subject. Printed in bold is the strongest differential regularisation strength before data residuals started to increase.

#### Inspection of streamline weight/delta coefficient distributions

We further evaluated the distributions of streamline weights (absolute optimisation) and delta coefficients (differential optimisation) at different regularisation strengths, determining the threshold at derived solutions are no longer contain a disproportionally large number of extreme streamline weights or delta coefficients.

The results of this analysis are shown in Supplementary Fig. 4 and 5 and can be summarised as follows:

- For absolute optimisation we found regularisation to take effect at a regularisation strength of approximately $10^{-1}$.
- For differential optimisation we found regularisation to take effect at a regularisation strength of approximately $10^{-1}$ (phantom) and $10^{-2}$ (in vivo) if the delta function was used, and approximately $10^{-2}$ (phantom) and $10^{-3}$ (in vivo) if the dual inverse barrier function was used.

Note that Supplementary Fig. 5 and 6 demonstrate the distribution of streamline weights/delta-coefficients only for streamline-based regularisation. Not shown are results for fixel and group-based regularisation, which yielded comparable results.

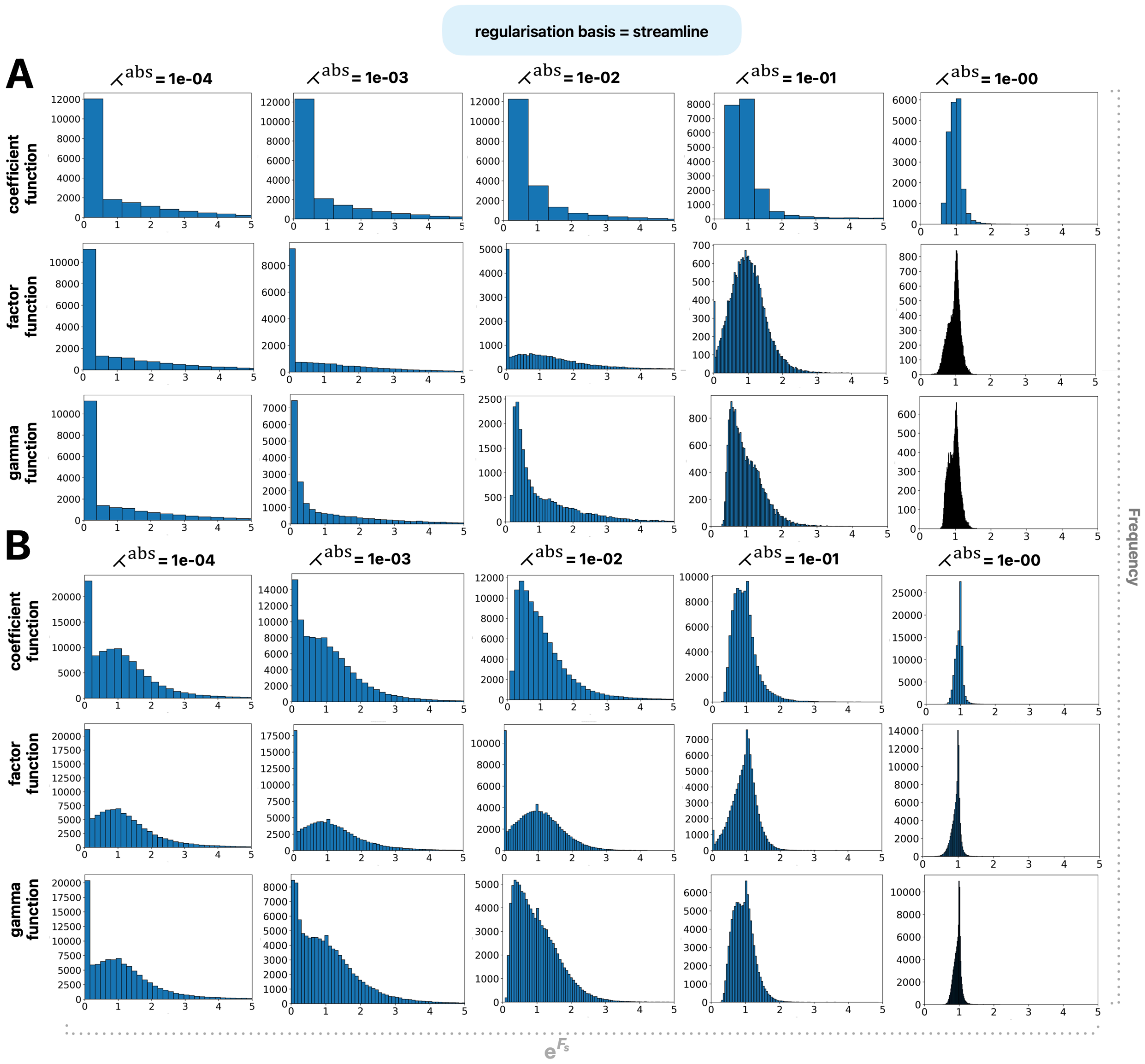


**Supplementary Fig. 5 – Distributions of absolute weights**. Shown are the distributions of streamline weights ($e^{F_s}$) for different regularisation functions (rows) at different regularisation strengths (columns). Panel A shows results derived in a synthetic dMRI phantom and panel B shows results derived in exemplar in vivo subject. Presented results were computed on a per streamline basis; results computed on a per-fixel and per-group basis (not shown) yielded highly similar results to those computed on a per-streamline basis $e^{F_s}$ = streamline weight, ʎ$^{abs}$ = regularisation strength for absolute optimisation.

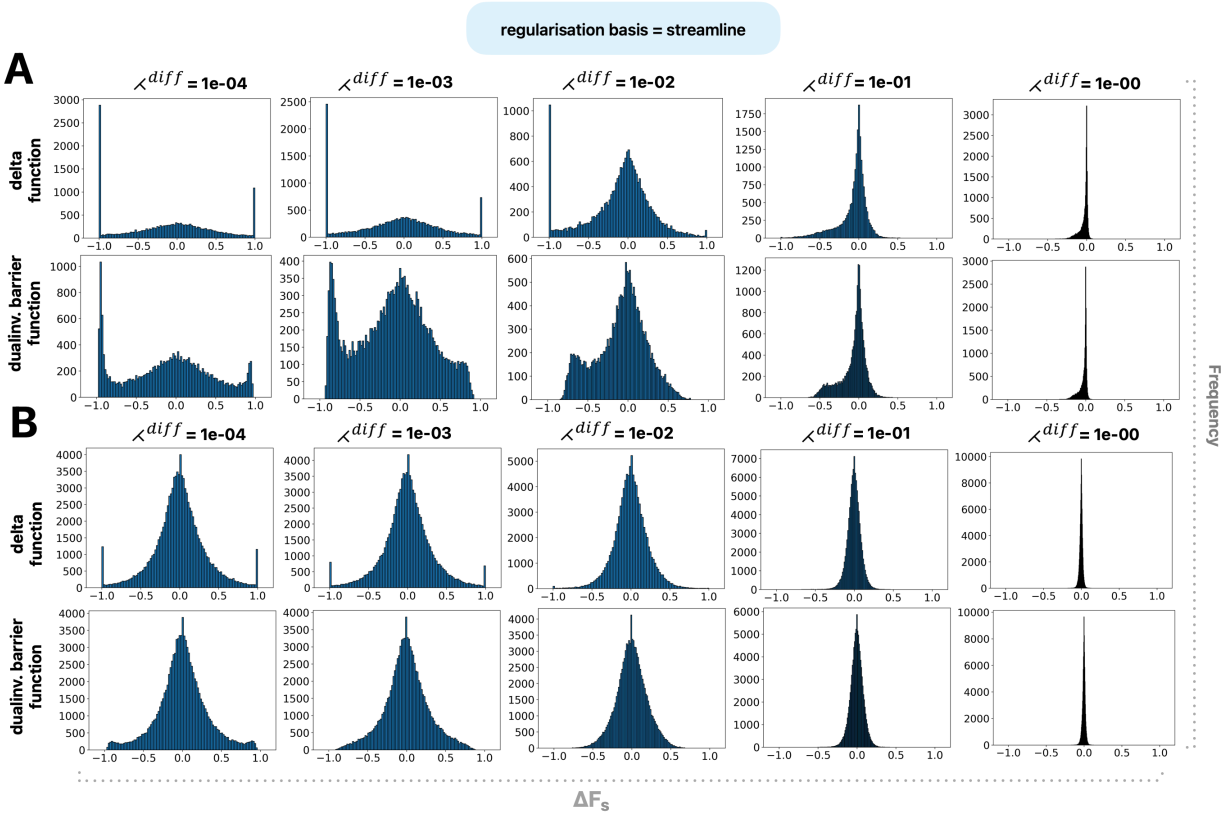


**Supplementary Fig. 6 – Distribution of delta coefficients**. Shown are the distributions of delta coefficients ($\Delta F_s$) for different regularisation functions (rows) at different regularisation strengths (columns). Panel A shows results derived in a synthetic dMRI phantom and panel B shows results derived in exemplar in vivo subject. Presented results were computed on a per streamline basis; results computed on a per-fixel and per-group basis (not shown) yielded highly similar results to those computed on a per-streamline basis. $\Delta F_s$ = delta coefficient, $\lambda^{diff}$ = regularisation strength for differential optimisation.

#### Error analysis

In our phantom with known ground truth, we evaluated edge-wise errors in absolute connectivity and connectivity differences at different regularisation strengths.

Absolute errors in structural connectivity were estimated as:

$$\varepsilon_{i,j}^{abs} = \left|\varphi \cdot SC_{ij} - FC_{ij}\right|$$

, where $\varphi$ is a global scaling factor to compare connectivity estimates with the underlying fibre count (see main manuscript), $SC_{i,j}$ is the estimated structural connectivity between parcels *i* and *j,* and $FC_{i,j}$ is the number of underlying ground truth fibres between parcels *i* and *j.*

Errors in absolute structural connectivity were then aggregated within categories, namely true bundles, false-positive bundles and all bundles. Within each of these categories we computed the mean absolute error:

$$MAE_{cat}^{abs} = \frac{1}{N}\sum_{i=0}^{N}\sum_{j=i}^{N}\varepsilon_{i,j}^{abs}$$

, where $MAE_{cat}^{abs}$ is the mean absolute error within bundle category *cat,* and $\varepsilon_{i,j}^{abs}$ the error in absolute structural connectivity estimated between parcels *i* and *j.*

Errors in edge-wise structural connectivity differences were estimated as $\varepsilon_{i,j}^{diff}$ (see main manuscript) and aggregated within categories, namely true bundles with effect, true bundles without effect, false-positives and all bundles. Within each of these categories we computed the mean absolute error:

$$MAE_{cat}^{diff} = \frac{1}{N}\sum_{i=0}^{N}\sum_{j=i}^{N}\varepsilon_{i,j}^{diff}$$

, where $MAE_{cat}^{diff}$ is the mean absolute error within bundle category *cat,* and $\varepsilon_{i,j}^{diff}$ the error in the estimated structural connectivity difference between parcels *i* and *j.*

The results of this analysis are presented in Supplementary Figure 7 (absolute optimisation) and Supplementary Figure 8 (differential optimisation), and can be summarised as follows:

- Analysis of $MAE_{cat}^{abs}$ indicated the optimal regularisation strength to lie at $10^0$ for streamline-based regularisation, $10^1$ for fixel-based regularisation, and $10^3$ for group-based regularisation
- Analysis of $MAE_{cat}^{diff}$ indicated the optimal regularisation strength to consistently lie at $10^{-1}$.

The error curves for specific bundle categories showed the largest $MAE_{cat}^{abs}$ within the "true bundles" and the largest $MAE_{cat}^{diff}$ within the "true bundles with effect" categories, which can be well explained based on how errors manifest in this analysis:

- For absolute optimisation, errors occur when densities are incorrectly distributed between "true bundles" or misattributed to "false positive" bundles. Since both of those errors will manifest in the "true bundles" category, and the "false positive" category encompasses a much larger set of bundles across which errors can occur, it seems plausible that $MAE_{cat}^{abs}$ is larger for the "true bundles" category.
- For differential optimisation, errors occur when longitudinal effects are incorrectly distributed between "true bundles with effect" or misattributed to "false positive" or "true bundles without effect"; this is likely to occur when multiple bundles traverse extended regions where local fibre density differences were observed, making it difficult for the algorithm to reliably attribute effects to the correct bundles. Since all of these errors will manifest in "true bundles with effect", and "false positive" and "true bundles without effect" encompass a much larger set of bundles across which errors can occur, it seems plausible that $MAE_{cat}^{diff}$ is largest for the "true bundles with effect" category.

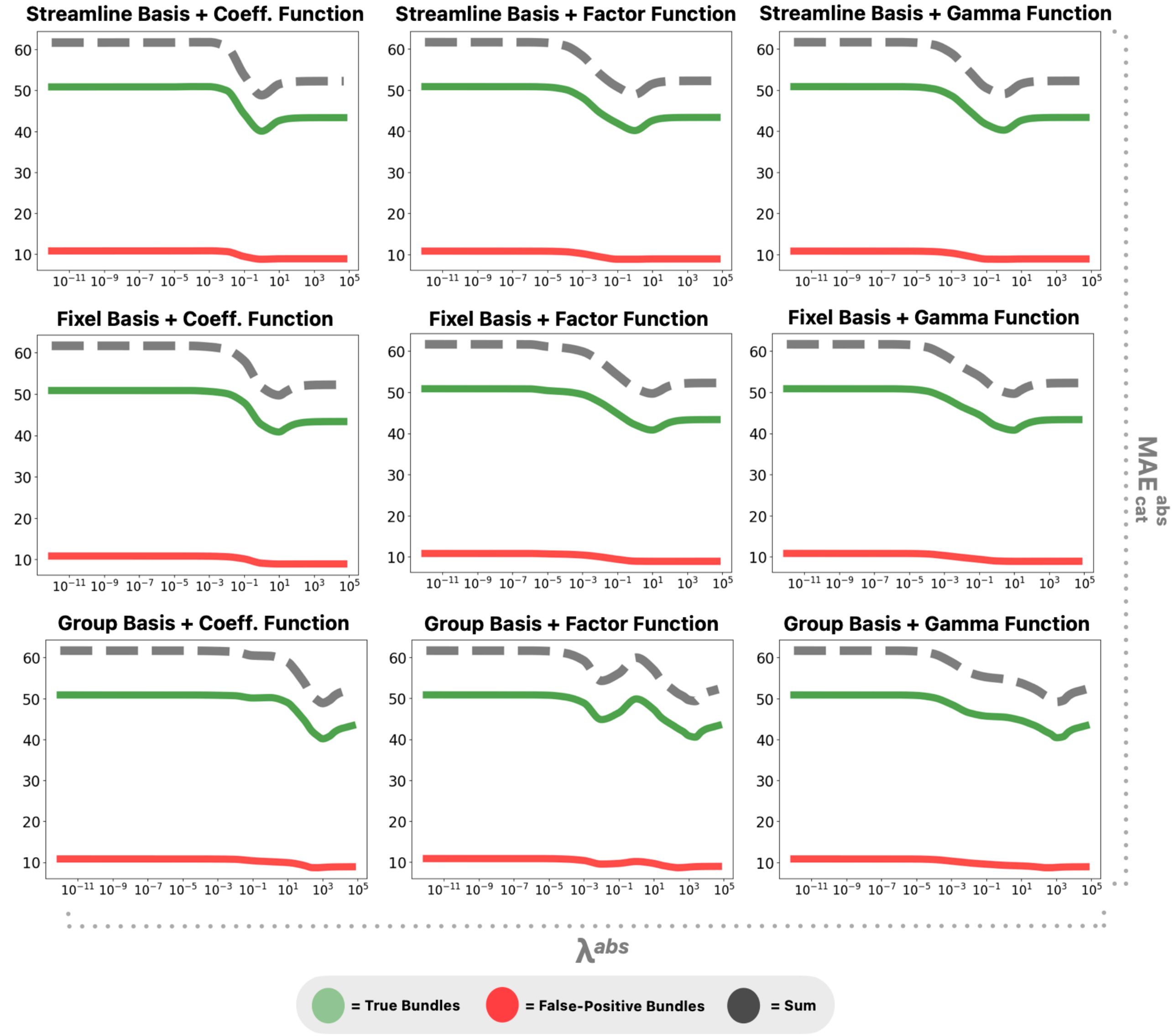


**Supplementary Fig. 7 – Absolute reconstruction errors at different regularisation strengths.** Shown is the mean absolute reconstruction error ($MAE_{cat}^{abs}$) within different bundle categories (true positives, false-positives) at different regularisation strengths; grey lines represent the sum of means across categories. The panels are organised so that rows show different regularisation bases and columns different regularisation functions, with each individual panel corresponding to one specific basis/function combination. $\lambda^{abs}$ = regularisation strength for absolute optimisation.

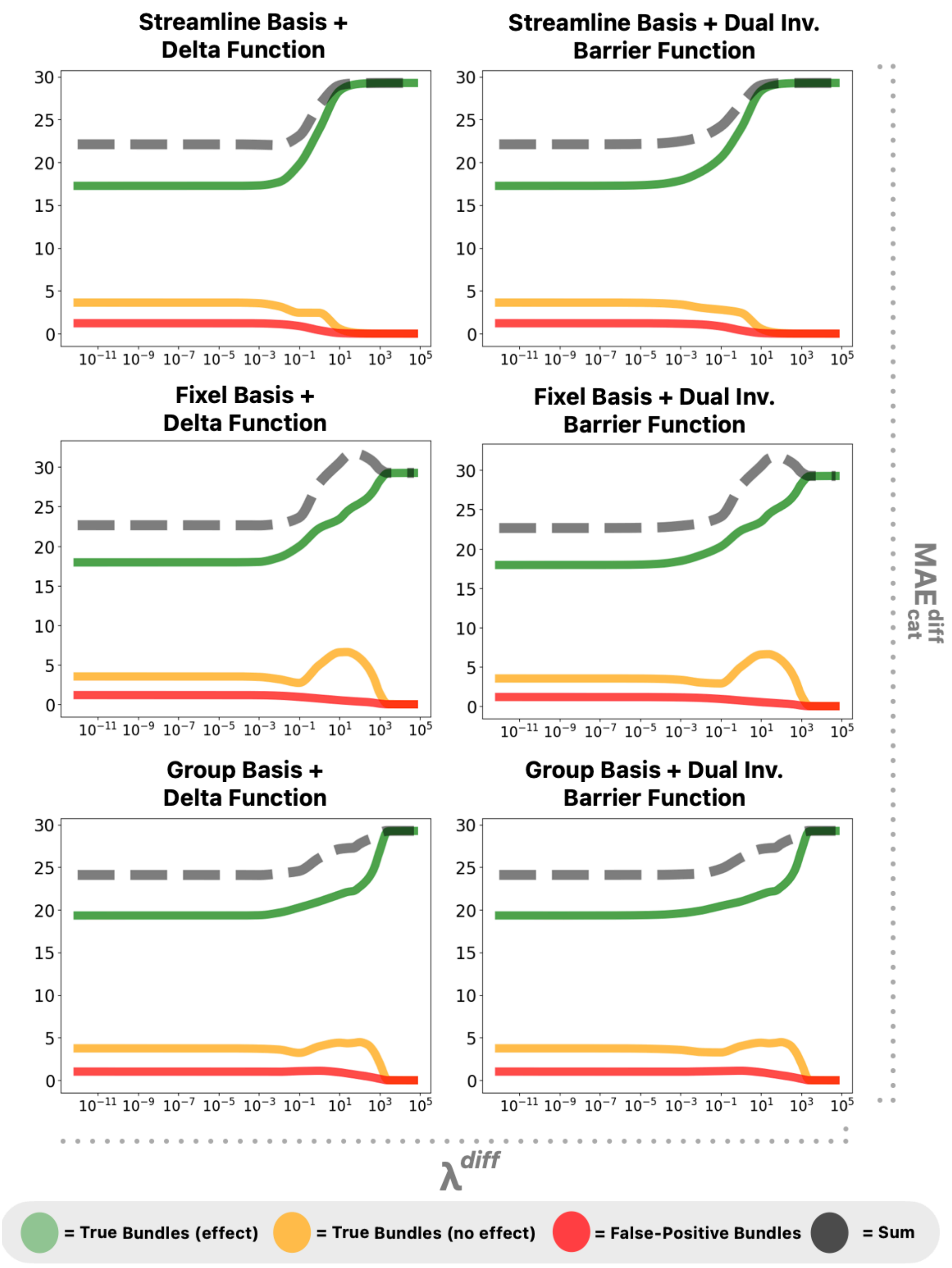


**Supplementary Fig. 8 – Differential reconstruction error at different regularisation strengths.** Shown is the mean longitudinal reconstruction error ($MAE_{cat}^{diff}$) within different bundle categories (true positives with, true positives without effect, false-positives) at different regularisation strengths; grey lines represent the sum of means across categories. The panels are organised so that rows show different regularisation bases and columns different regularisation functions, with each individual panel corresponding to one specific basis/function combination. $\lambda^{diff}$ = regularisation strength for differential optimisation.

#### Synthesis of results

We conducted several analyses to choose an appropriate regularisation strength for absolute and differential optimisation, including L-curve analyses, inspection of the distributions of streamline weights/delta coefficients and analyses of bundle-wise errors against a known ground truth.

The results can be summarised as follows:

- L-curve analyses of absolute and differential regularisation functions suggested the ideal regularisation strength to lie at approximately $10^{-4}$ to $10^{-3}$.
- Visual inspection of the distributions of optimised values however indicated solutions at these strengths to still be dominated by few individual streamlines. The frequency of extreme values substantially decreased at regularisation strengths of $10^{-1}$ and above.
- Error analyses indicated the optimal regularisation strength to lie at approximately $10^{0}$ to $10^{3}$ for absolute regularisation (depending on the chosen basis) and $10^{-1}$ for differential regularisation. While it is unclear how well the specific thresholds generalise to human *in vivo* data, the analysis provides evidence to support regularisation strengths above those considered "ideal" in the L-curve analyses.

In light of these findings, we believe that an absolute and differential regularisation strength of $10^{-1}$ provides a sensible choice for most use cases. Given that errors were comparable between the different regularisation bases, there does not seem to be a clear benefit of the more complex fixel and group-wise over simple streamline-wise regularisation. Hence for this article, we chose to perform absolute and differential regularisation on a per-streamline basis, using the gamma (absolute) and dual inverse barrier (differential) functions at a regularisation strength of $10^{-1}$.

#### Derivation of synthetic dMRI data

For *in silicio* validation, we synthesised dMRI images of three longitudinal models, all based on the previously published DiSCo3 phantom(Rafael-Patino et al., 2021). Longitudinal effects were simulated by modulating the fibre count between timepoints. From these ground truth fibres, artificial dMRI images were synthesised using Fiberfox(Neher et al., 2014).

Simulation settings were as follows:

- Image Settings:
  - Image Dimensions: 40×40×40 voxels
  - Image Spacing: 1×1×1 mm
  - Diffusion sampling scheme: b=0 (4), b=1000 (90), b=2000 (90), b=3000 (90), with gradient directions as provided in the original DiSCo3 publication(Rafael-Patino et al., 2021)
- Intra-axonal compartment: Stick Model
  - T1: 974 ms
  - T2: 70 ms
  - Axial Diffusivity: 0.0017 $mm^2/s$
- Extra-axonal compartment: Ball Model
  - T1-Relaxation: 4658 ms
  - T2-Relaxation: 2200 ms
  - Diffusivity: 0.003 $mm^2/s$
- Noise and other artefacts: none simulated at this stage (noise added post simulation for improved control of signal-to-noise ratio)

Using these settings, we found Fiberfox to produce dMRI images with unrealistic signal drops between shells that did not resemble human data. Further investigation revealed this to arise due to the relatively low white matter volume fraction of the DiSCo3 phantom (approximately 0.3), which is well below that expected in human white matter (approximately 0.7(Zhang et al., 2012)). This low fibre density, in combination with the fibre radius that was used within the forward model (per default automatically determined by Fiberfox), led to the synthesisation of unrealistic dMRI images. We rectified this by empirically determining an appropriate fibre radius for the forward model, so that final dMRI images had signal drops that resembled human data (see Supplementary Fig. 9). To these images, we added Rician noise at a signal-to-noise ratio of 20(Garyfallidis et al., 2014).

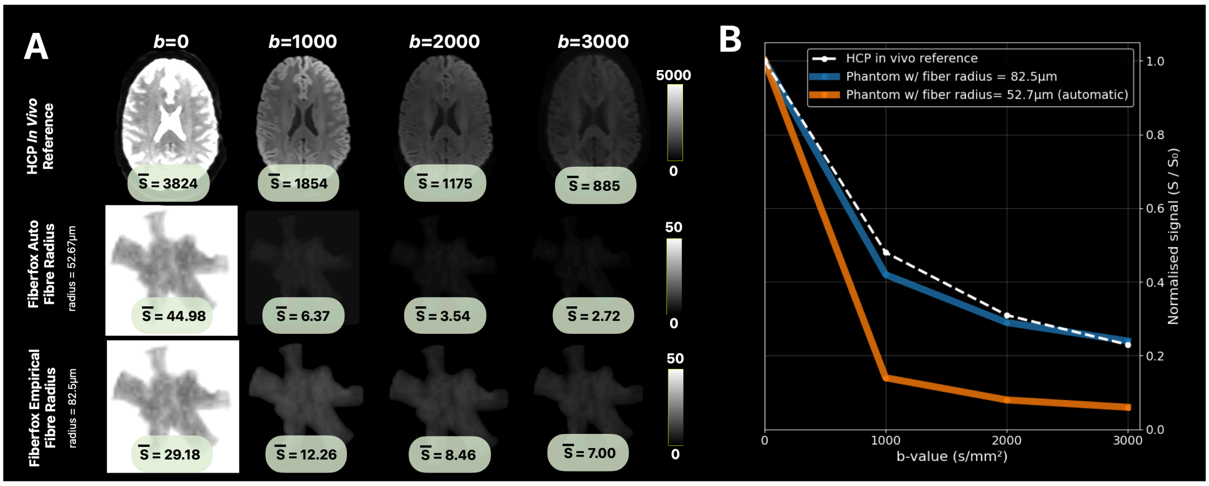


**Supplementary Fig. 9 – Synthesising dMRI images with realistic signal drops between shells. Panel A**: the first row shows expected drops in the mean signal intensity across the different shells in an exemplar subject of the HCP Scan-Rescan cohort. The second row shows the Fiberfox output image that was synthesised using our initial settings, producing signal drops that did not resemble human *in vivo* data. We found this issue to arise due to the relatively low density of fibres within the DiSCo3 phantom, which was rectified by determining an appropriate fibre radius used by the forward model, with outputs using this custom radius shown in the third row. **Panel B**: Signal drops (normalised to mean $b$=0 image) as a function of b-value. The white line shows the expected signal drops of a reference in vivo subject; orange and blue lines show the signal drops of the initial Fiberfox output and one using the empirically determined fibre radius. Abbreviations: HCP = Human Connectome Project, $\bar{S}$ = mean signal intensity of brain voxels.

##### Cohort Characteristics HCP Scan-Rescan Data

The HCP Scan-Rescan dataset comprises healthy young adults between 22 and 35 years old and were scanned approximately six months apart.

Scans were acquired on a Siemens 3T Connectom Skyra(Fan et al., 2022) scanner with a 32-channel head coil:

- T1w sequence: 3D MPRAGE, TR: 2400ms, TE: 2.14ms, TI: 1000ms, flip angle: 8°, field of view: 224×224×224 mm, voxel size: 0.7mm$^3$, parallel imaging: GRAPPA factor 2, acquisition time: 7:40.
- dMRI sequence: spin-echo EPI with multiband acceleration, TR: 5520ms, TE: 89.5ms, flip angles: excitation 78°, refocusing 160°, field-of-view: 210×180 mm, matrix: 168×144, voxel size: 1.25mm$^3$, slices: 111, multiband factor: 3, echo spacing: 0.78ms, phase partial Fourier: 6/8, b-values: 1000, 2000, and 3000 s/mm, directions: 90 diffusion directions per shell + 18 b=0 volumes.

##### Cohort Characteristics Developing Children Data

Developing Children were aged between six and eight years and scanned approximately one year apart.

Scans were acquired on a 3T Philips Achieva scanner with a 32-channel head coil:

- T1w sequence: 3D T1TFE, TR: 8.93 ms, TE: 4.61 ms, flip angle: 8°, voxel size: 1 × 1 × 1 mm, imaging matrix: 256 × 256, acquisition time: 4 min 25 s.
- dMRI sequence: spin-echo EPI, TR: 2900 ms, TE: 79 ms, voxel size: 2.1 × 2.1 × 2.2 mm, 40 diffusion directions at b = 1000 s/mm$^2$ and 56 diffusion directions at b = 2000 s/mm$^2$ + 16 b=0 volumes, acquisition time: 6 min 41 s.

##### Cohort Characteristics Temporal Lobe Surgery Data

Temporal Lobe Surgery was performed in adults between 18 and 68 years for treatment of epilepsy with scans acquired approximately 1 (IQR 2.4) month before and 18 (IQR 23.5) months after surgery. Patients were either treated with selective amygdalohippocampectomy (n=36), anterior temporal lobectomy (n=13) or laser interstitial thermal therapy (n=3).

Temporal Lobe Surgery scans were acquired on a 3T Philips Achieva scanner with a 32-channel head coil.

- T1w sequence: 3D T1TFE, TR: 9 ms, TE: 4.61 ms, flip angle: 8°, voxel size: 1 × 1 × 1 mm, imaging matrix: 256 × 256, CS-SENSE acceleration factor: 3, acquisition time: 2 min 20s.
- dMRI sequence: spin-echo EPI, TR: 7100 ms, TE: 64 ms, voxel size: 2.5 × 2.5 × 2.5 mm, 92 diffusion directions at b = 1600 s/mm² + one b=0 volume, acquisition time: 11min 16s.

#### Additional considerations for image processing of surgical data

The presence of gross resections introduces several complexities for unbiased image processing, as a significant fraction of white matter being present only in the preoperative timepoint violates the framework's assumption of fixed gross anatomy over time. We hence made additional adjustments to account for the absence of resected tissue in the postoperative image data. Resections cavities required for these processing steps were segmented using an automated workflow described elsewhere(Pruckner et al., 2025).

Additional adjustments for surgical pipelines included:

- Within-subject template generation: Averaging timepoints will result in implausible average intensities within the area of tissue resection. Hence, we derived anatomically intact within-subject templates by imputing preoperative tissue intensities into the surgical resection cavity(Pruckner et al., 2025).
- Pipeline 3 (symmetric optimisation): weights of those streamlines within the unbiased template tractogram that intersected the resection cavity were set to zero before initialising postoperative optimisation, excluding them from further refinement.
- Pipeline 4 (differential optimisation): delta coefficients of the unbiased template tractogram that intersected the resection cavity were set to -1 (=100% connectivity loss from first to second time point) and remained fixed throughout differential optimisation.
- Parcellation: portions of brain regions that were resected were removed from the atlas segmentations, restricting the analysis to only the non-resected portions of partially resected brain regions(Pruckner et al., 2025).

### Supplementary Results

#### Unbiased reconstruction results in less errors in false-positive pathways

To investigate whether the lower errors in quantified structural connectivity changes using unbiased compared to cross-sectional pipelines (see Fig. 7) was predominantly due to one specific category of bundles, we repeated the statistical analysis separately for true bundles with effect, true bundles without effect and false-positive bundles. Specifically, we tested for significant differences in bundle-wise longitudinal error ($\varepsilon_{i,j}^{diff}$) (see Methods, Validation in *in silicio* dMRI phantoms) between Pipelines 1-4 in the central lesion phantom for each of these bundle categories.

The analysis yielded the following results (see Supplementary Fig. 10):

- True bundles with effect showed no significant differences in $\varepsilon_{i,j}^{diff}$ between reconstruction pipelines.
- True bundles without effect showed no significant differences in $\varepsilon_{i,j}^{diff}$ between reconstruction pipelines.
- False-positive bundles showed significantly lower $\varepsilon_{i,j}^{diff}$ estimated through unbiased reconstruction Pipelines 3 & 4, as compared to cross-sectional reconstruction Pipelines 1 & 2. No significant differences were found between Pipelines 1 & 2 or between Pipelines 3 & 4.

This analysis provides evidence to support that improvements of unbiased reconstruction pipelines are primarily driven by suppression of quantified effects within false-positive pathways.

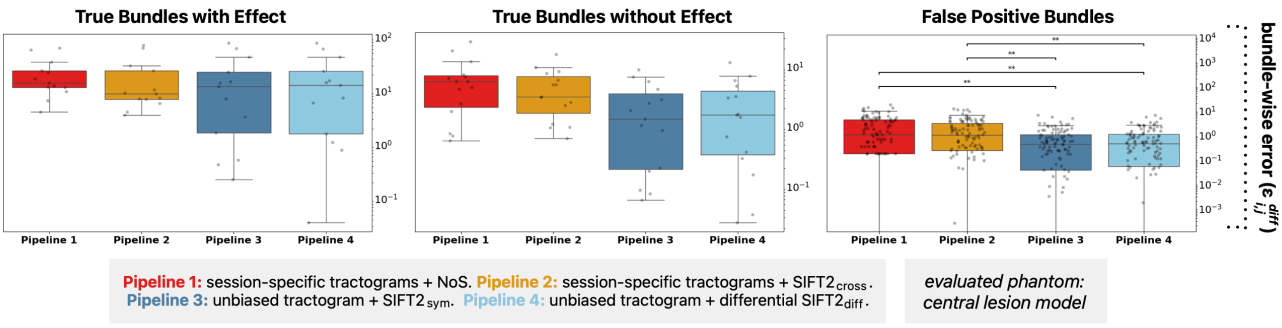


**Supplementary Fig. 10 – Post-hoc error analysis for specific bundle-categories.** Shown are the pipeline-wise errors in estimated longitudinal connectivity changes in the central lesion model. Panels show the bundle-wise error, plotting the fibre count error for each individual bundle; percentages are expressed in respect to each bundle's fibre count at the first timepoint for true positives bundles or the mean fibre count across all bundles at the first timepoint for false-positive bundles. Asterisks (**) indicate statistically significant differences ($p<0.01$). cross = cross-sectional, diff = differential, SIFT = Spherical Deconvolution Informed Filtering of Tractograms, sym = symmetric.

#### Unbiased optimisation reduces methodological imprecision

TFNBS yielded results for unbiased pipelines that were both more extensive and more concordant with biological expectation (see Fig. 8), which we hypothesised to be due to a decrease in edge-wise imprecision. To investigate whether or not this was indeed the case, for each cohort and pipeline, we computed for every edge the across-subject standard deviation of ΔSC. Within each cohort, we then tested for statistical differences in the mean edge-wise standard deviation across pipelines through one-way ANOVA omnibus testing; if significant, differences between pairs of pipelines were assessed using Tukey's Honestly Significant Difference post-hoc tests. Results were considered significant at $p<0.05$ after adjustment for multiple comparisons.

The results of this analysis are shown in Supplementary Fig. 11 and can be summarised as follows:

- Across all cohorts, Pipelines 3 & 4 led to significantly lower edge-wise standard deviations of ΔSC compared to Pipelines 1 & 2.
- No significant differences in edge-wise standard deviations of ΔSC were found between Pipelines 1 & 2 or between Pipelines 3 & 4.

In summary, these findings demonstrate that unbiased reconstruction pipelines substantially decreases edge-wise imprecision, providing a tangible explanation for their improved statistical sensitivity to biological effects in in vivo cohorts.

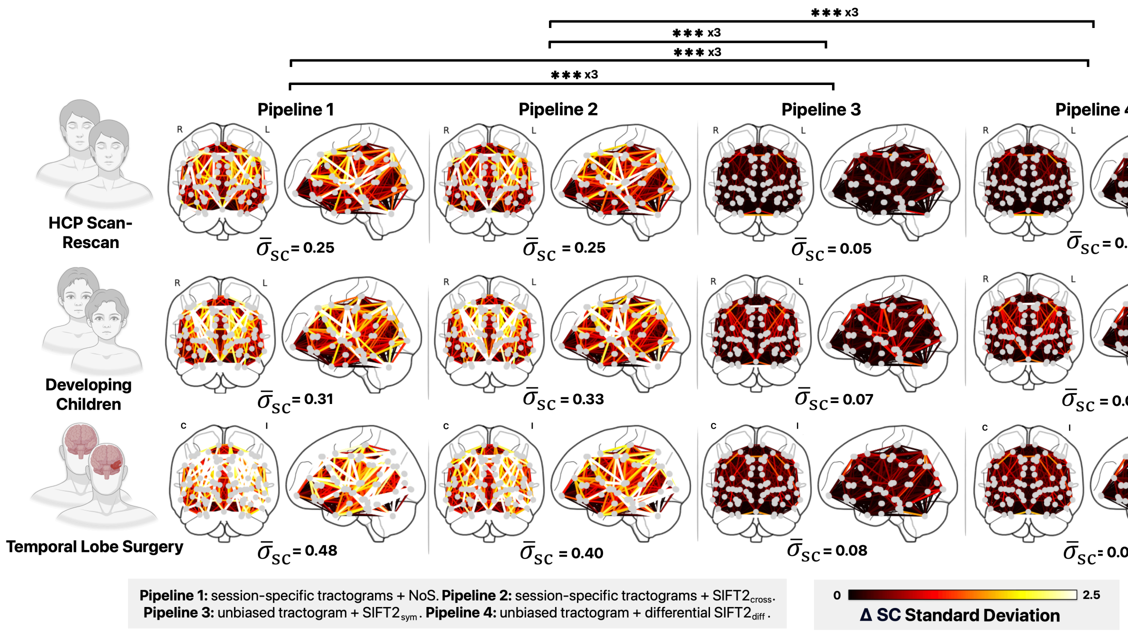


**Supplementary Fig. 11 - Edge-wise imprecision across pipelines.** Connectome plots visualise the edge-wise standard deviation of estimated structural connectivity changes (ΔSC), with annotations showing the mean standard deviation across all edges ($\bar{\sigma}_{sc}$). Asterisks (***x3) indicate statistically significant differences ($p<0.001$) between pipelines across all three cohorts. C/I = contralateral/ipsilateral, cross = cross-sectional, diff = differential, L/R = left/right, NoS = Number of Streamlines, SIFT = Spherical Deconvolution Informed Filtering of Tractograms, sym = symmetric.

#### References


Fan, Q., Eichner, C., Afzali, M., Mueller, L., Tax, C. M. W., Davids, M., Mahmutovic, M., Keil, B., Bilgic, B., Setsompop, K., Lee, H.-H., Tian, Q., Maffei, C., Ramos-Llordén, G., Nummenmaa, A., Witzel, T., Yendiki, A., Song, Y.-Q., Huang, C.-C., … Huang, S. Y. (2022). Mapping the human connectome using diffusion MRI at 300 mT/m gradient strength: Methodological advances and scientific impact. *NeuroImage*, *254*, 118958. https://doi.org/10.1016/j.neuroimage.2022.118958

Garyfallidis, E., Brett, M., Amirbekian, B., Rokem, A., Van Der Walt, S., Descoteaux, M., & Nimmo-Smith, I. (2014). Dipy, a library for the analysis of diffusion MRI data. *Frontiers in Neuroinformatics*, *Volume 8-2014*. https://www.frontiersin.org/journals/neuroinformatics/articles/10.3389/fninf.2014.00008

Neher, P. F., Laun, F. B., Stieltjes, B., & Maier-Hein, K. H. (2014). Fiberfox: Facilitating the creation of realistic white matter software phantoms. *Magnetic Resonance in Medicine*, *72*(5), 1460–1470. https://doi.org/10.1002/mrm.25045

Pruckner, P., Mito, R., Vaughan, D., Jackson, G., Fischmeister, F., Heinz-Nenning, K., Berger, M., Pataraia, E., Baumgartner, C., Dorfer, C., Rössler, K., Czech, T., Kasprian, G., Bonelli, S., & Smith, R. (2025). Beyond the Resection: Surgical White Matter Disruption Structurally Alters Non-Resected Brain Anatomy. *bioRxiv*. https://doi.org/10.1101/2025.03.19.643850

Rafael-Patino, J., Girard, G., Truffet, R., Pizzolato, M., Caruyer, E., & Thiran, J.-P. (2021). The diffusion-simulated connectivity (DiSCo) dataset. *Data in Brief*, *38*, 107429. https://doi.org/10.1016/j.dib.2021.107429

Smith, R. E., Tournier, J.-D., Calamante, F., & Connelly, A. (2015). SIFT2: Enabling dense quantitative assessment of brain white matter connectivity using streamlines tractography. *NeuroImage*, *119*, 338–351. https://doi.org/10.1016/j.neuroimage.2015.06.092

Zhang, H., Schneider, T., Wheeler-Kingshott, C. A., & Alexander, D. C. (2012). NODDI: Practical in vivo neurite orientation dispersion and density imaging of the human brain. *NeuroImage*, *61*(4), 1000–1016. https://doi.org/10.1016/j.neuroimage.2012.03.072
